# A single-molecule reporter of membrane-proximal actin detects rapid remodeling upon B cell receptor clustering

**DOI:** 10.64898/2026.04.23.720115

**Authors:** Adam Decker, Sarah Veatch

## Abstract

Membrane-proximal (MP) actin represents the subset of cortical f-actin localized within 10 nm of the plasma membrane. Here, we describe a family of single-molecule MP actin (SM-MPAct) probes that diffuse within the plasma membrane and are transiently immobilized through binding to f-actin, enabling the localization of MP actin in both time and space. These probes quantify aspects of MP actin structure, dynamics, and remodeling by analyzing probe positions using a combined single-particle tracking and correlation-function approach. This is demonstrated using chemical and physical perturbations of actin and actin-binding proteins, and by interrogating MP actin organization and dynamics in early B cell receptor (BCR) activation. Upon crosslinking of the IgM BCR, MP actin transiently remodels to increase the size of actin corals, facilitating the efficient assembly of BCR clusters and the local accumulation of MP actin. Notably, analogous remodeling is not detected in measurements using total f-actin probes, indicating that SM-MPAct is uniquely sensitive to the f-actin pool that regulates signaling processes at the plasma membrane.

**STATEMENT OF SIGNIFICANCE:** The actin cortex provides mechanical stability to the plasma membrane and contributes to the organization and dynamics of plasma membrane components. This report presents single-molecule probes and analytical methods to characterize the density, mesh size, motion, and turnover dynamics of the portion of the actin mesh in direct contact with the plasma membrane, enabling quantitative studies of actin remodeling in plasma membrane processes. This general framework is demonstrated through quantification of actin remodeling during early B cell receptor signaling and could be applied to a broad range of cell processes.

## INTRODUCTION

Actin is coupled to the plasma membrane and serves a variety of structural and functional roles. Interactions between actin and the membrane are regulated by a network of actin-binding proteins and actin assembly factors (recently reviewed in (1, 2)). This coupling is crucial for a wide range of cellular functions, including cell motility, surface adhesion, maintenance of cell junctions, endocytosis, phagocytosis, and synapse formation. To accomplish these functions, cells tune the biophysical properties of both actin and the plasma membrane, coordinating the assembly of specialized structures at the cell surface. The attachment of actin to the plasma membrane also regulates the organization and mobility of plasma membrane components (3, 4). While it is clear that f-actin organization and dynamics are critical for function at the plasma membrane, and vice versa, the detailed mechanisms underlying this co-regulation remain poorly understood. One common approach is to perturb actin and actin-binding proteins through chemical perturbations; however, these treatments typically dramatically alter actin organization and dynamics (5), making it difficult to infer f-actin’s specific functional contributions in the unperturbed state.

The organization and dynamics of f-actin can be directly visualized using fluorescent actin-binding probes. Common approaches for labeling actin employ transient actin-binding domains derived from full-length proteins, like lifeact or ftractin (6, 7), or small actin-binding toxins, such as phalloidin (8–10). These actin-binding molecules are conjugated to fluorescent dyes or proteins, visualizing the distribution of actin in cells. Actin probes are often imaged using TIRF microscopy, which limits the illumination depth to approximately 200 nm to observe the actin cortex. Single-molecule microscopy methods, such as PALM, STORM, and PAINT, have also been applied to resolve the nanoscale structure and organization of actin in both fixed (11–13) and live cells (14–18). These methods often require modifications of actin-binding probes with photoswitchable, blinking, or transient-binding fluorophores. One challenge in applying these actin imaging tools is that conventional f-actin probes label all cellular actin, making it difficult to isolate the pool of actin in direct contact with the membrane. Even when using TIRF microscopy methods, the imaging depth (∼200 nm) is still an order of magnitude greater than distances most relevant to direct and effective interactions between actin and plasma membrane components (<20nm) (19).

To better isolate the signal from membrane-proximal (MP) actin, f-actin-binding domains have been modified by adding a membrane anchor, restricting binding within approximately 10 nm of the plasma membrane (20–25). A subset of these probes use low-affinity f-actin-binding domains that transiently bind MP actin (20–22) and are collectively referred to as MPAct (20). These probes have been used with advanced fluorescence microscopy techniques to demonstrate how actin affects the mobility of plasma membrane components (22–25).

Here, we present a family of single-molecule MPAct (SM-MPAct) probes, along with analytical methods that together quantify the organization and dynamics of MP actin over a broad range of spatial and temporal scales. Probes transiently immobilize when bound to MP actin, enabling its localization in both time and space. We interrogate MP actin organization and dynamics primarily in the CH27 B cell line, and demonstrate the effects of temperature and several inhibitors of f-actin and associated proteins. We also quantify MP actin remodeling and colocalization with B-cell receptor (BCR) clusters during BCR activation and find that SM-MPAct better isolates the MP actin population that assembles in response to activation than probes that detect total f-actin. More generally, this work provides experimental and analytical approaches for investigating MP actin in live cells, enabling direct measurements of MP actin remodeling and its interactions with individual plasma membrane proteins.

## RESULTS AND DISCUSSION

### Immobile SM-MPAct is bound to actin

SM-MPAct is schematically depicted in **Figure 1A**. In SM-MPAct, the conventional fluorophore is replaced with one that can photo-switch between fluorescent and non-fluorescent states, facilitating the imaging of single-molecule motions using live-cell single-molecule localization microscopy. The CH27 B cell shown in **Figure 1B** was imaged for 7 min under total internal reflection (TIR) illumination at room temperature. This cell is expressing a SM-MPAct probe consisting of the ftractin f-actin binding domain, the mEos3.2 photo-switchable protein fluorophore, and a membrane anchor consisting of a series of basic amino acids proximal to a geranylgeranyl (GG) motif (Ftractin-mEos3.2-GG). A movie showing a series of raw images of diffusing probes for this cell is provided in **Movie 1**.

**Figure 1:**
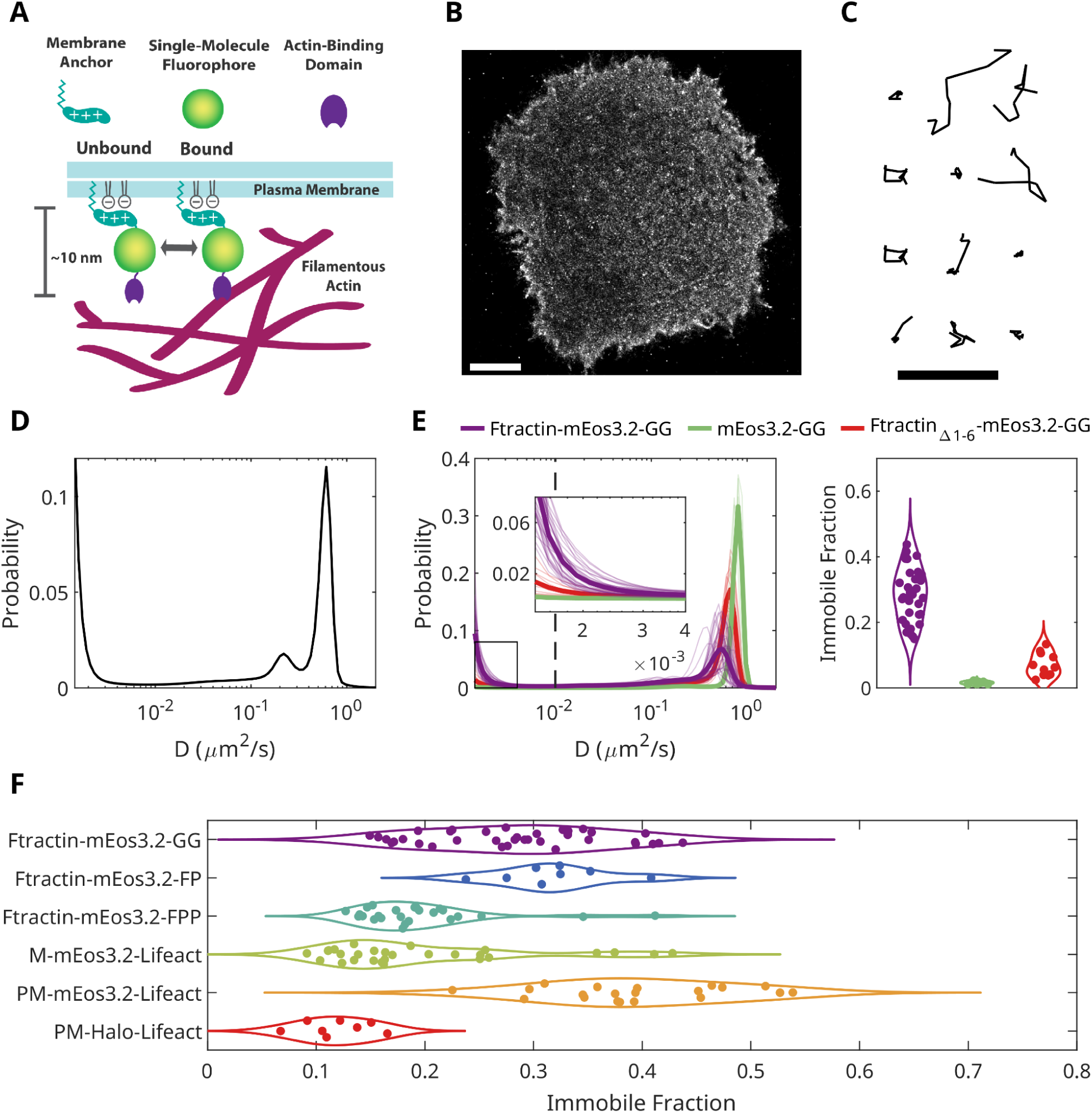
SM-MPAct immobilizes when bound to actin. **(A)** Schematic of a SM-MPAct probe that localizes to the plasma membrane and transiently binds to MP actin. **(B)** Reconstruction of Fractin-mEos3.2-GG localizations acquired through imaging a live CH27 cell for 7 minutes (15000 image frames). Scale bar is 5 µm. **(C)** Representative long trajectories (10 frames each) of Ftractin-mEos3.2-GG from the cell in panel B. Scale bar is 1 µm. **(D)** SASPT posterior diffusion coefficient distribution for the cell shown in panel B. **(E)** (right) Posterior distributions for cells expressing Ftractin-mEos3.2-GG (N=37), mEos3.2-GG (N=6), or FtractinΔ1-6-mEos3.2-GG (N=11). Curves from individual cells are drawn as thin lines and the average drawn as thick lines. (left) The fraction of immobile segments determined from integrating the posterior distribution below a cutoff of D = 10^−2^ µm^2^ s^−1^. **(F)** Fraction of immobile SM-MPAct across cells expressing Ftractin-mEos3.2-GG (N=37), Ftractin-mEos3.2-FP (N=8), Ftractin-mEos3.2-FPP (N=25), M-mEos3.2-Lifeact (N=29), PM-mEos3.2-Lifeact (N=20), or PM-Halo-Lifeact (N=8).

Linking single-molecule localizations into trajectories enables characterization of probe mobility, and several representative trajectories are shown in **Figure 1C**. Visual impressions of both trajectories and motions of fluorescent molecules in Movie 1 suggest that this SM-MPAct probe can occupy both mobile and immobile states. This visual impression is validated through a state array single particle tracking (SASPT) analysis (26), which uses the ensemble of single-molecule trajectories to estimate the probability distribution that individual track segments have specific diffusion coefficients (**Figure 1D**) (see methods). The distribution of diffusion coefficients (D) of Ftractin-mEos3.2-GG in this single cell contains three distinct peaks: two populations with peaks near D = 0.6 and 0.15 µm^2^ s^−1^, and a third population peaked at the lowest diffusion value available to the SASPT algorithm. From the distribution of diffusion coefficients, we interpret that there are at least three distinct diffusive states for SM-MPAct, two mobile states and one immobile state.

The three mobility states are assigned using variants of this SM-MPAct probe. **Figure 1D** also shows the average distribution of diffusion coefficients for multiple cells expressing the full-length Ftractin-mEos3.2-GG alongside average distributions of cells expressing a truncated version lacking the entire ftractin domain (mEos3.2-GG) or a version where the N-terminal 6 amino acids of the ftractin domain are removed (FtractinΔ1-6-mEos3.2-GG), reducing its f-actin binding affinity (20). Movies depicting raw images of different cells expressing the 3 probes are provided in **Movie 2**. Only one mobile state is detected from the ensemble of mEos3.2-GG trajectories, while 2 mobile states are identified in the ensemble of FtractinΔ1-6-mEos3.2-GG trajectories. Based on these observations, we assign the fast mobile state of Ftractin-mEos3.2-GG to freely diffusing membrane probes and the immobile state to probes bound to f-actin. The slower mobile state detected in both Ftractin-mEos3.2-GG and FtractinΔ1-6-mEos3.2-GG likely reflects ftractin domain interactions with cellular components that are distinct from its direct actin binding capacity. The presence of the ftractin domain or its truncated variant also slows and broadens the faster mobile state peak compared to the mEos3.2-GG probe, again suggesting that it mediates alternate interactions, likely at time-scales on or faster than the frame-rate of these measurements (22 ms).

Since the immobile and mobile states are well separated, the fraction of immobile segments is extracted by integrating probability distributions below a cutoff of D = 10^−2^ µm^2^ s^−1^ (**Figure 1E**). As expected, the fraction of immobile segments is small for both mEos3.2-GG (1.6 ± 0.2%) and FtractinΔ1-6-mEos3.2-GG (7 ± 1%) and is larger for Ftractin-mEos3.2-GG (28 ± 1%). There is significant cell-to-cell variation in the fraction of immobile segments, even though the fraction for a single cell is well specified, indicating that there is variation in the density of cortical actin accessible to this SM-MPAct probe across cells.

The modular nature of SM-MPAct allows for a range of fluorophores, f-actin, and membrane binding motifs. **Figure 1F** reports the immobile fraction of a panel of probes expressed in CH27 cells and imaged at room temperature. These include three probes with the same mEos3.2 fluorophore and ftractin actin binding domain, but distinct membrane binding motifs, the GG motif described above, in addition to ones taken from N-Ras (FP) and H-Ras (FPP). These also include 3 probes with a lifeact f-actin binding domain. Two are conjugated to the mEos3.2 fluorophore and membrane binding motifs from c-Src (M) and Lyn (PM), and the last is conjugated to a halotag and the PM binding motif and visualized with a haloligand conjugated to janeliafluor635b (JF635b) (27). These probes report a wide range of immobile fractions, indicating that their f-actin binding domains have different accessibility to MP-actin. While one goal of the measurements in Figure 1F was to explore how the mode of membrane anchoring affects its accessibility to MP actin, Alphafold3 (28) structures predict structural differences beyond those engineered into the constructs that may also contribute to the results **(Figure S1)**. In particular, differences in the secondary structures of linker sequences differ across probes. In addition, structures for PM-Halo-Lifeact predict the existence of direct interactions between the halotag and lifeact domains, consistent with the low immobile fractions detected for this probe. Overall, this indicates that careful probe design would be required to draw conclusions when swapping specific motifs.

### The immobile fraction of SM-MPAct reports MP actin density for a given SM-MPAct probe

The physical attachment of actin to the plasma membrane is highly regulated across many biochemical networks and is mediated by a collection of membrane-to-cortex adaptor proteins and actin assembly factors (2, 29, 30). Cells actively tune the density of cortical actin to regulate biochemical and mechanical homeostasis, maintaining a gradient in the distribution of actin perpendicular to the cell surface (19, 31, 32). **Figure 2** explores how probe mobility is affected by chemical and physical perturbations of actin and actin-binding proteins.

**Figure 2.**
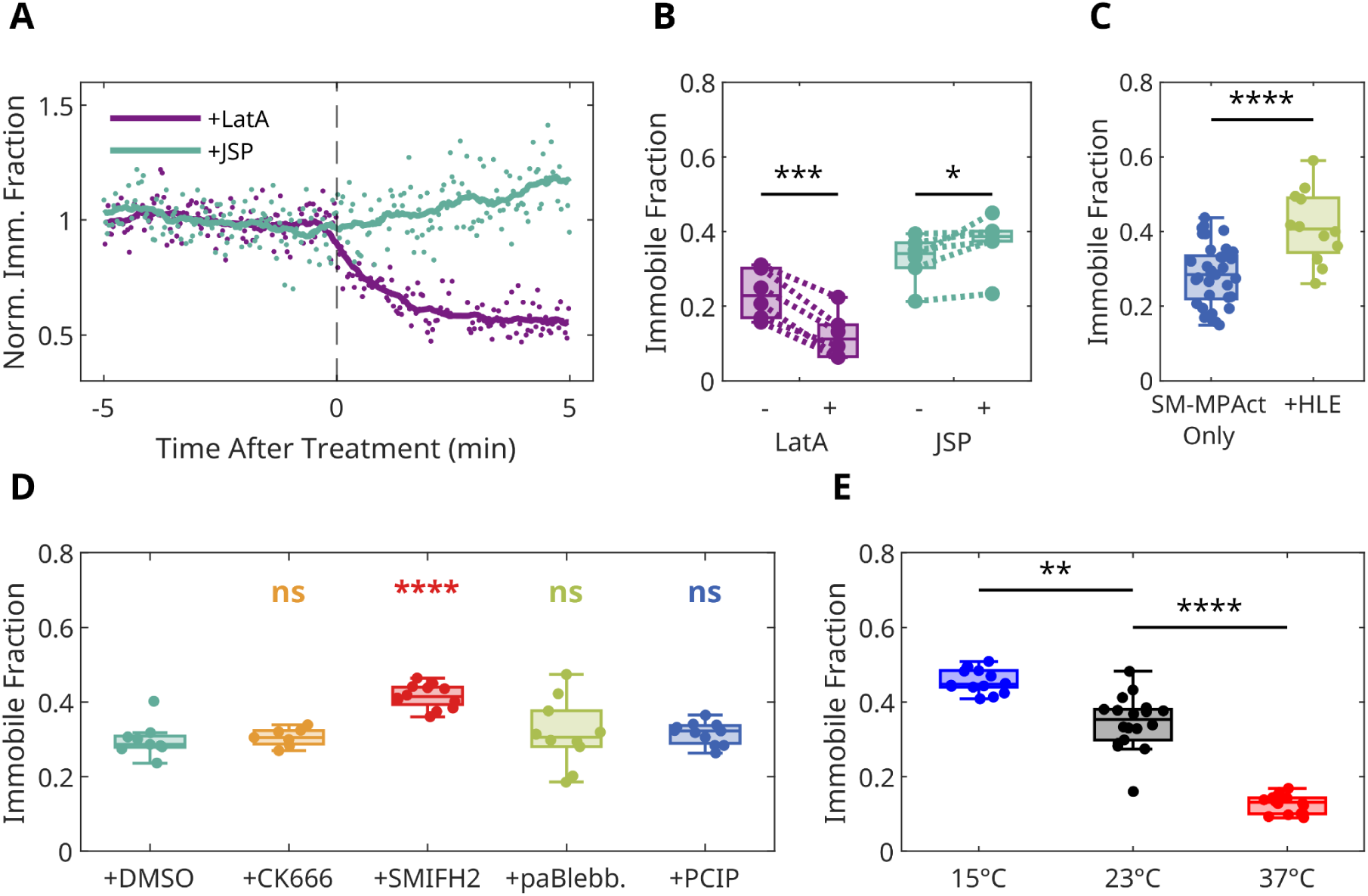
The immobile fraction of a given SM-MPAct probe quantifies MP actin density. **(A)** Time-resolved immobile fraction from single cells expressing Ftractin-mEos3.2-GG treated with either 0.5 µM LatA or 1 µM JSP at t=0, normalized to the average immobile fraction before treatment. **(B)** Immobile fractions compared across populations of cells expressing Ftractin-mEos3.2-GG before and after treatment with LatA (N=6) or JSP (N=6). **(C)** Immobile fractions for cells expressing Ftractin-mEos3.2-GG (N=37 replotted from Figure 1E) or coexpressing Ftractin-mEos3.2-GG and HLE (N=12). **(D)** Immobile fractions for cells expressing PM-mEos3.2-Lifeact treated with 2% (v/v) DMSO (N=9), 100 μM CK666 (N=7), 40 μM SMIFH2 (N=12), 25 μM paBlebb (N=10), or 5 μM PCIP (N=11). **(E)** Immobile fractions for cells expressing PM-mEos3.2-Lifeact observed at 15°C (N=14), 23°C (N=18), or 37°C (N=16). Significance in A is determined using a 1-tailed test, comparing the same cells before and after treatment. Significance in C, D, and E is determined using a 2-tailed t-test, comparing the perturbed population to the SM-MPAct-only, DMSO, or 23°C population, respectively. Symbols for p-values from significance testing represent *p≤0.05,**p≤0.01,***p≤0.001,****p≤0.0001, or ns for p>0.05

The f-actin inhibitor latrunculin A (LatA) binds to monomeric actin, inhibiting f-actin growth and reducing the concentration of f-actin within the cell, including at the plasma membrane (33, 34). Exposing a single CH27 cell expressing Ftractin-mEos3.2-GG to 0.5 µM of LatA produces a time-dependent drop in the fraction of immobile segments **(Figure 2A)**. The f-actin binding drug Jasplakinolide (JSP) stabilizes f-actin and inhibits its dynamics (35, 36). Exposing a single cell to 1 µM of JSP has a modest effect on the fraction of immobile segments. These observations are robust across cell populations, as indicated in **Figure 2B**. We detect a systematic increase in the fraction of immobile segments in cells coexpressing a constitutively active chimera of the ezrin protein embedded within the plasma membrane via a transmembrane helix (HLE) (37) **(Figure 2C)**. This Ezrin variant binds to f-actin with high affinity and stabilizes interactions between the plasma membrane and cortical actin, making f-actin more accessible to SM-MPAct. Each of these perturbations to MP actin has the expected outcome and supports that shifts in the immobile fraction quantify changes in the density of MP actin.

We additionally exposed cells to a range of inhibitors of f-actin-associated proteins to probe their impact on MP actin density in CH27 cells expressing the SM-MPAct probe PM-mEos3.2-Lifeact **(Figure 2D)**. CH27 cells treated with 100 μM of the Arp2/3 inhibitor CK666 did not show a significant change in the immobile fraction, indicating that the average density of MP actin does not significantly change under this condition. In contrast, inhibiting formins with 40 μM SMIFH2 increases the density of MP actin, consistent with previous reports (38). We also disrupted myosin I or myosin II motor activity by treating with either 25 μM para-amino blebbistatin (paBlebb) or 5 μM pentachloropseudilin (PCIP), respectively. In both cases, MP actin density was not significantly impacted. The immobile fraction of SM-MPAct is also affected by changes in temperature **(Figure 2E)**, with fewer immobile molecules at 37°C and more at 15°C, which we attribute to temperature-dependent changes in the binding kinetics between lifeact and f-actin.

### Spatio-temporal correlation functions report on MP actin motion and remodeling

The motions of SM-MPAct probes can also be characterized by tabulating the autocorrelation function, g(r, *τ*) (39–41). g(r,*τ*) reports on the density of localizations separated by a distance r and time-interval *τ* from the average localization of the same type, normalized by the density expected if localizations were dispersed randomly in space and time. Since most fluorophores are detected over multiple sequential frames, g(r, *τ*) has a large amplitude at short *τ* with a spatial extent that reflects the distribution of displacements from single fluorophores (**Figure 3A**). We further evaluate g(r,*τ*) by calculating the integrated intensity, I(*τ*), which quantifies the magnitude of correlations at each *τ*. I(*τ*) decays sharply because single molecules are only visualized for a few sequential frames, and fitting I(*τ*) to an exponential decay (see Methods) extracts the characteristic fluorophore off-time, *τ*_fluor_, of 0.12 ± 0.01 s (∼ 5 frames) for this cell. The shape of g(r) at each *τ* is well described by a superposition of two Gaussian functions with different associated mean-squared displacements (MSDs). Examining how these MSDs evolve over *τ* indicates distinct populations of mobile and immobile probes (**Figure S2**), in good agreement with results from single-molecule tracking. We extract the lifetime of actin binding by tabulating the fraction of correlated segments within the immobile peak, which is *τ*_off_ = 0.25 ± 0.02 s for this cell, corresponding to roughly 12 frames.

**Figure 3.**
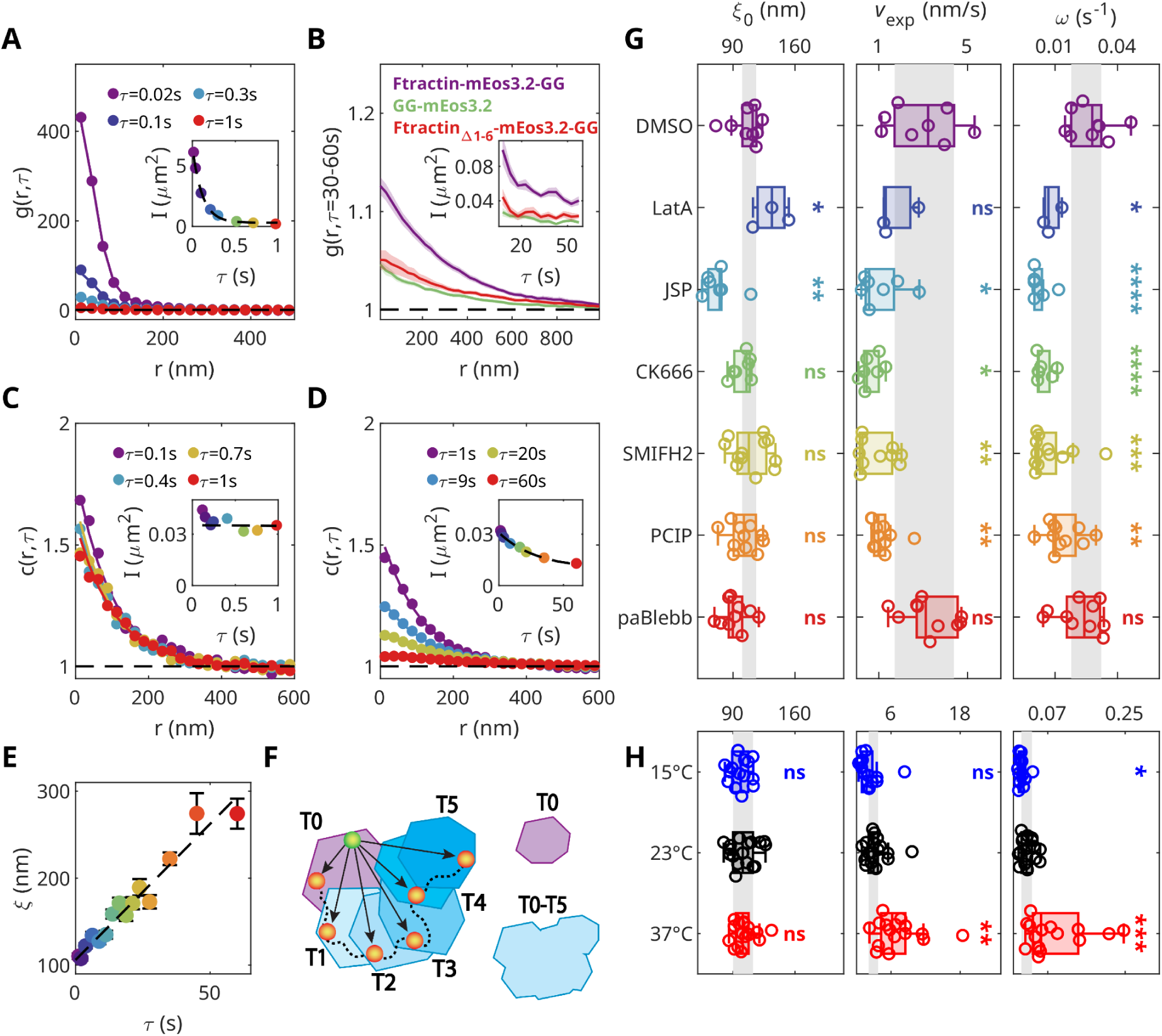
Spatial-temporal correlation functions report on average MP actin structure, lateral motion, and turnover. **(A)** The autocorrelation, g(r,*τ*), of Ftractin-mEos3.2-GG in a CH27 cell plotted at several distinct time-intervals, *τ*, and fit to a sum of two Gaussian functions as described in Methods. The inset shows the normalized integrated intensity, I(*τ*), fit to Aexp(-*τ*/*τ*_fluor_)+C. **(B)** g(r,*τ* = 30-60 s) averaged over multiple cells expressing Ftractin-mEos3.2-GG (N=37), GG-mEos3.2 (N=6), or FtractinΔ1-6-mEos3.2-GG (N=11). The inset displays the average I(*τ*) of g(r,*τ*). Error bars represent the standard error of the mean. **(C, D)** The crosscorrelation, c(r,*τ*), between PM-mEos3.2-Lifeact and PM-Halo(JF635b)-Lifeact in a single CH27 cell, measured across varying short (C) and long (D) *τ* ranges and fit to Aexp(-r/ξ(*τ*))+C. Insets show I(*τ*) fit to Aexp(-⍵*τ*)+C. **(E)** ξ(*τ*) values obtained from the curves in panel D, fit to ξ_0_+v_exp_*τ*. **(F)** Cartoon depicting the expansion of MP actin mesh size with *τ* due to the translation of actin corals. **(G, H)** ξ_0_, v_exp_, and ⍵ evaluated across the same cells as presented in Figure 2D,E treated with actin inhibitors (G) or imaged at different temperatures (H). Significance tests in F and G are the results of a 2-tailed t-test comparing perturbed cells with the DMSO vehicle control or cells observed at 23°C, respectively. Shaded regions represent the range between the first and third quartiles of the DMSO control or 23°C populations. Symbols for p-values from significance testing represent *p≤0.05, **p≤0.01, ***p≤0.001, ****p≤0.0001, or ns for p>0.05

While the amplitude of g(r, *τ*) is greatly reduced for time intervals longer than the blinking time of individual fluorophores, correlations persist at long *τ* as evident in plots of g(r) tabulated for *τ* between 30 and 60 s (**Figure 3B)**. Similar long *τ* correlations are diminished for probes that do not bind MP-actin, suggesting that this elevated signal arises from correlated motions of MP-actin itself. To confirm, we conducted a two-color measurement in which two distinguishable MPAct probes were observed simultaneously over time, then tabulated the crosscorrelation function c(r, *τ*). Unlike the autocorrelation, c(r, *τ*) does not report on single-molecule motions because the detection of different colored molecules is uncorrelated. Correlations between different molecules arise from their shared interactions with MP-actin, and as a consequence, the shape and amplitude of c(r) are largely constant for *τ* < 1 s (**Figure 3C**). The magnitude of c(r) instead decays over time-scales of seconds to minutes (**Figure 3D**), reflecting the dynamics of MP actin. Consistent with this interpretation, similar long *τ* correlations are detected when SM-MPAct is imaged alongside soluble Halo-Lifeact, but are greatly reduced in measurements where one or both probes lack f-actin binding domains **(Figure S3)**. The integrated intensity, I(*τ*), of the two color SM-MPAct c(r,*τ*) gradually decreases, reflecting MP actin disassembly and exchange with the cytoplasmic f-actin pool. We fit the I(*τ*) to an exponential decay as described in Methods to extract a remodeling rate, ⍵, for MP actin, which is found to be ⍵= 0.034 ± 0.007 s^−1^ for this cell. This value indicates that MP actin remodeling occurs over tens of seconds, consistent with previous measurements for total cortical f-actin remodeling at room temperature (42).

Spatial correlations at constant *τ* decay at increasing separation distances, and it is convenient to parameterize trends in c(r,*τ*) by fitting to an exponential decay, where ξ is the decay length. At short *τ*, c(r) closely resembles g(r) curves obtained when total f-actin is imaged in chemically fixed CH27 cells under TIR excitation, consistent with both observations reflecting the steady state structure of cortical f-actin **(Figure S4)**. In both cases, ξ is close to 100 nm, which we interpret to be the characteristic size of f-actin corrals at steady state, in good agreement with previous measurements (43). In live cells, ξ(*τ*) increases linearly with *τ* **(Figure 3E)**. One interpretation of this finding is that ξ(*τ*) increases due to translations of the actin meshwork, effectively expanding the corral size when comparing localizations across time **(Figure 3F)**. We further parameterize these findings by fitting ξ(*τ*) to a linear trend, extracting the intercept, ξ_0_ = 106 ± 3 nm, and slope, v_exp_ = 3.1 ± 0.2 nm s^−1^, which represent the steady state coral size and expansion speed, respectively. In this example, the expansion speed is well specified and small, taking roughly 30s for the average corral size to double from 100 to 200 nm.

The impact of drugs that perturb actin or actin-binding proteins is displayed in **Figure 3G**. As expected, both LatA and JSP arrest the temporal evolution of c(r). LatA also increases the steady-state corral size, while JSP modestly reduces it, consistent with previous reports (38) and our observations of altered MP actin density with these drugs reported in Figure 2. Pretreatment with CK666 or SMIFH2, which inhibit Arp2/3 or formins, respectively, also arrests MP actin dynamics, as evidenced by slower remodeling rates and expansion speeds compared with a vehicle control. Myosin I inhibition with PCIP also arrested MP actin dynamics; however, myosin II inhibition with paBlebb did not significantly alter remodeling rates or the expansion speeds. The different observations for myosin I vs. myosin II inhibition are consistent with myosin I being more membrane-proximal than myosin II (44, 45).

Lastly, we compared MP actin dynamics within CH27 cells imaged at different temperatures (**Figure 3H)**. As expected, dynamics are significantly impacted by temperature, as reflected in changes in both remodeling times and expansion speeds. The expansion speed increases by about 3-fold, and the remodeling rate increases by about 4-fold for cells observed at 37 °C compared to cells observed at 23 °C. Cells observed at 15 °C exhibit a modest but significant ⅔-fold decrease in the expansion speed and remodeling rate, having a less dramatic effect on MP actin dynamics than JSP, CK666, SMIFH2, and PCIP. Interestingly, temperature does not affect the steady-state corral size. This analysis supports our previous conclusion in Figure 2E that temperature-dependent changes in the immobile fraction arise from different binding kinetics **(Figure S2)**. Overall, this approach can extract diverse information on the average structure and dynamics of MP actin in live cells.

### Images of immobile SM-MPAct visualize static actin structures

Results shown in Figures 1 and 2 analyzed the ensemble of trajectories from single cells to determine the fraction of immobile probes. It is also possible to use the SASPT framework to localize immobile segments, and therefore the positions of MP actin, in both time and space. This is accomplished by identifying the most probable diffusion coefficient over the entire trajectory segment based on the posterior distribution for that segment. When defined this way, track segments exhibit diffusion coefficients that again distribute into one immobile (actin-bound) and 2 mobile (unbound) states, as seen in the histogram of **Figure 4A** and representative segments from each population shown in **Figure 4B**. While this approach leads to the occasional miscategorization of immobile localizations, the vast majority are correctly identified since trajectory segments are short (∼3 frames on average) relative to the unbinding time for the SM-MPAct probes (∼10 frames).

**Figure 4:**
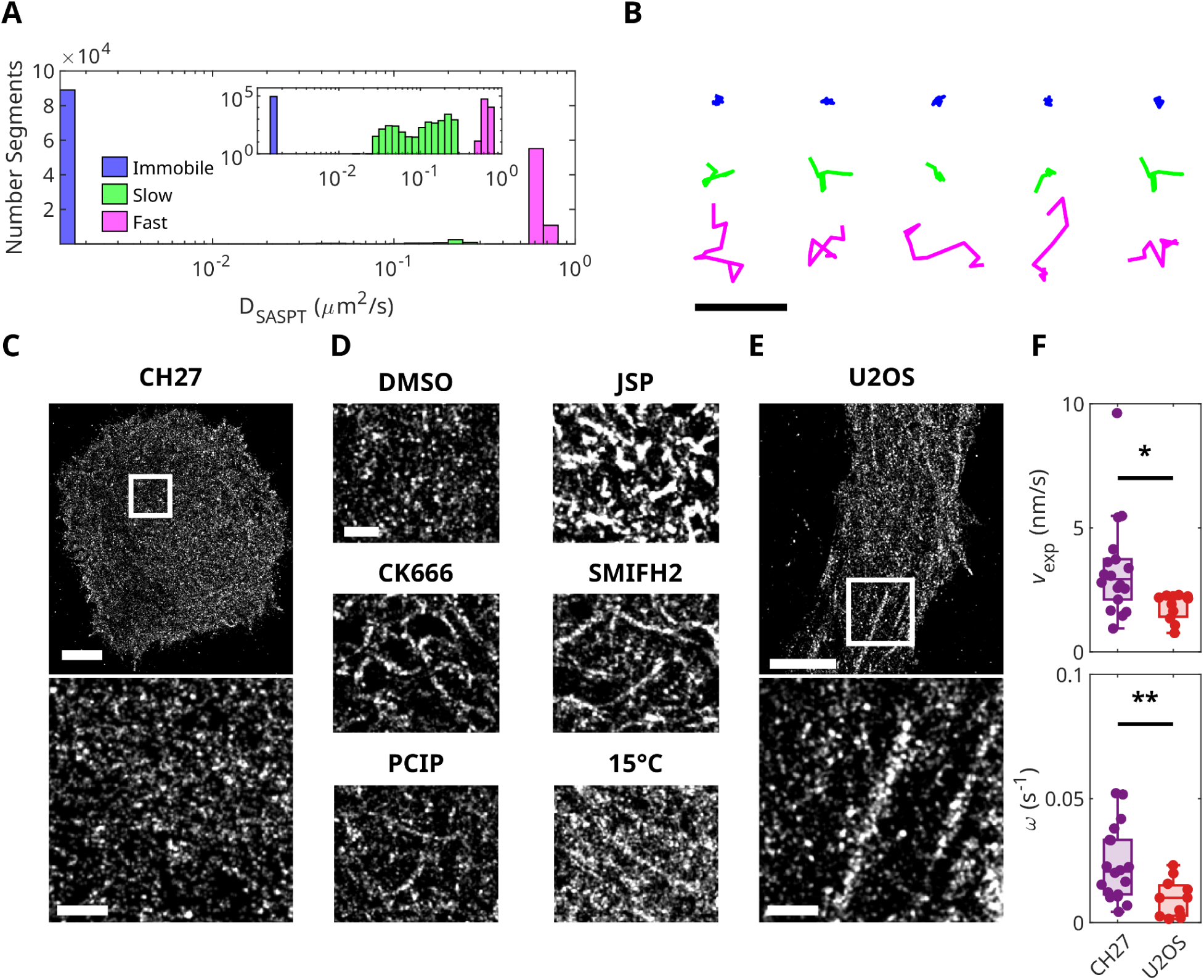
Immoble SM-MPAct visualizes static MP actin structures. **(A)** Histogram of diffusion coefficients (D) for trajectories from a CH27 cell expressing Ftractin-mEos3.2-GG determined as described in Methods. Trajectories are distributed into the three states, with D ≤ 0.01 μm² s^−1^ as the immobile state, 0.01 μm² s^−1^ < D < 0.3 μm² s^−1^ as the slow state, and D ≥ 0.3 μm² s^−1^ as the fast state. The inset shows the same histogram on a log-log axis. **(B)** Representative trajectories plotted and colored by their D with coloring as in panel B. Scale bar is 1μm. **(C)** Reconstructed image showing the positions of immobile trajectories from panel A. **(D)** Reconstructions showing subsets of CH27 cells evaluated in Figure 2B,C and Figure 3G,H expressing PM-mEos3.2-Lifeact treated with the indicated inhibitors or observed at 15 °C. **(E)** Reconstructed image of immoble PM-mEos3.2-Lifeact in a U2OS cell. **(F)** v_ξ_, and ⍵ evaluated from crosscorrelations of of PM-mEos3.2-Lifeact and PM-Halo-Lifeact in CH27 and U2OS cells. Significance is determined from a 2-tailed t-test. Scale bars in full-cell reconstructions and zoomed-in panels represent 5 μm and 1 μm, respectively. Symbols for p-values from significance testing represent *p≤0.05, **p≤0.01, ***p≤0.001, ****p≤0.0001, or ns for p>0.05

The localizations associated with immobile trajectories can be used to reconstruct images, as shown in **Figure 4C-E**, or movies, as shown in **Movie 3**. Reconstructed images of immobile segments acquired over 7 minutes in CH27 cells imaged at room temperature contain puncta distributed across the cell surface **(Figure 4C)**. This is expected, since 7 minutes is much longer than the remodeling time for MP actin determined for these same cells (∼40 seconds), and the measured expansion speed suggests that cortical filaments that remain at the cell surface translate several hundred nm over this time period. Reconstructing images over shorter time windows produces images that are poorly sampled in space, obscuring the visualization of cortical meshwork structures **(Movie 3)**. In contrast, robust cortical meshworks are detected in cells imaged under conditions with arrested MP actin dynamics **(Figure 4D)**. Under these conditions, the time-scales of remodeling and translation are long compared to the time required to assemble a well-sampled image.

**#Figure 4E** shows reconstructed images of MP actin in a representative U2OS cell. U2OS cells are an adherent cell type that exhibit extended and static structures with high MP actin densities visible in images reconstructed from 8 min of acquired localizations. In this example, stress fibers and focal adhesions are visible against a backdrop of diffuse, punctate structures, which likely represent more mobile cortical MP actin meshworks. In agreement with this interpretation, U2OS cells have a slower average expansion speed and remodeling rate compared to CH27 cells **(Figure 4F),** consistent with previously reported measurements for slow actin dynamics in cells with stress fibers (46). Overall, these results support the idea that MP actin is dynamic across a range of spatio-temporal scales, as visualized in reconstructions of immobile SM-MPAct and quantified by spatio-temporal correlations.

### MP actin is transiently depleted during BCR activation

Ligating the B-cell receptor (BCR) with a multivalent, soluble crosslinker leads to receptor clustering and activation, as well as the recruitment and activation of diverse membrane-bound and soluble proteins that propagate the cellular-level signaling response (40, 47, 48). Past work has demonstrated that BCR clustering, activation, and the mobility of clustered receptors are impacted by perturbations of f-actin organization and dynamics (49). Actin nucleating proteins are also recruited to BCR clusters downstream of receptor activation (50, 51). Here we apply SM-MPAct to further interrogate this system.

**Figure 5A** shows a representative reconstructed image of a CH27 B cell assembled from 5 min of localizations from BCR and Ftractin-mEos3.2-GG acquired both before and after BCR crosslinking with soluble streptavidin. BCR is visualized using a biotinylated anti-IgM antibody Fab (μ-chain specific) that is chemically conjugated with the Silicone Rhodamine (SiR) organic fluorophore and biotin. BCR is mobile in untreated cells, and both slows and becomes clustered soon after BCR crosslinking with streptavidin. BCR clusters are themselves dynamic, as visualized in a series of reconstructions rendered over shorter time windows **(Movie 4)**. In past work, we have characterized BCR clustering dynamics and signaling responses under the same stimulation and imaging conditions used in these measurements (39, 40, 52).

**Figure 5.**
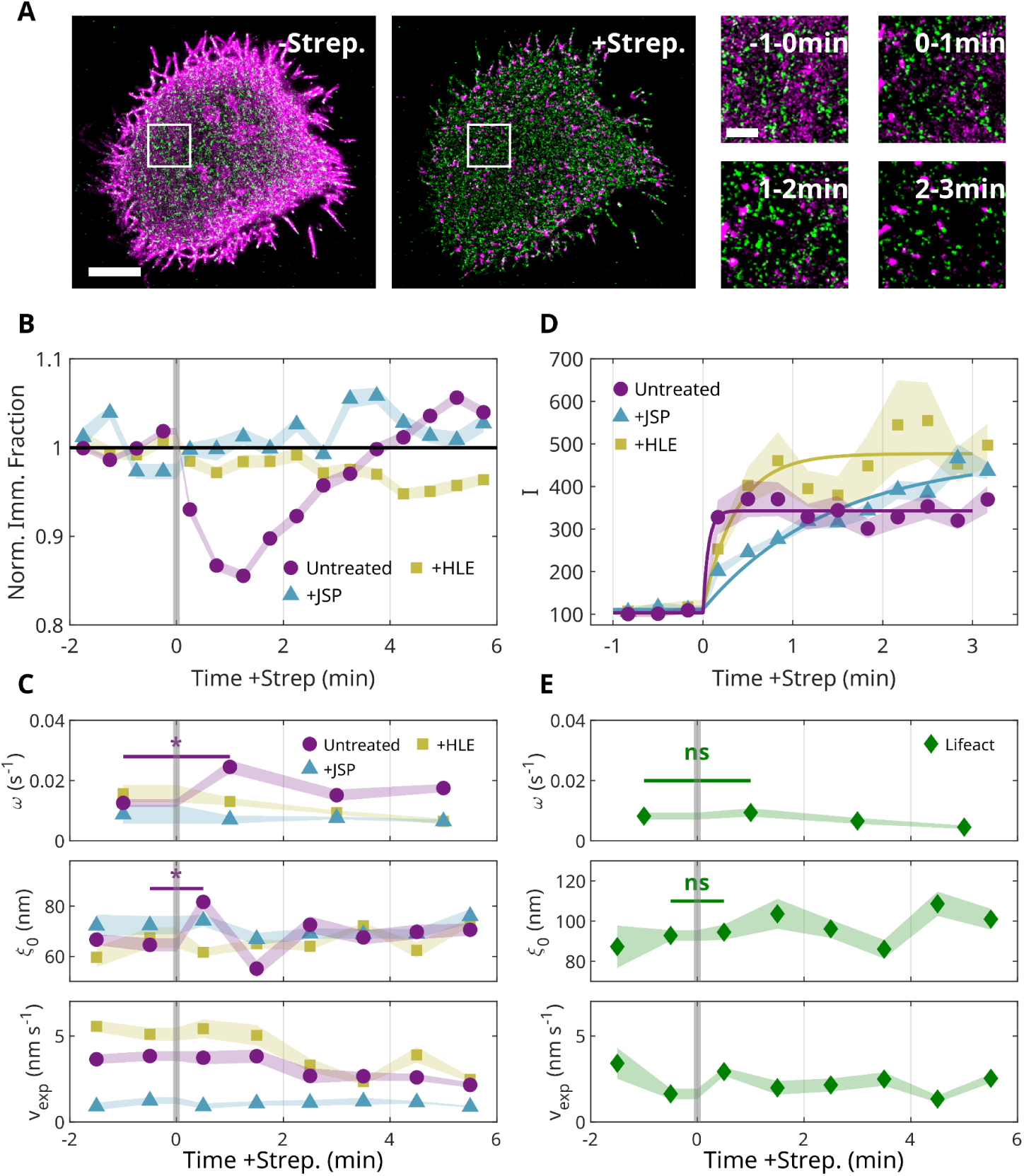
Transient depletion of MP actin supports rapid BCR clustering. **(A)** Reconstructed images showing BCR and immobile Ftractin-mEos3.2-GG rendered before and after BCR crosslinking with strepaviden. Images are generated from localizations acquired either 5 min before (left) or 1-6 min after strepaviden addition (middle), or as indicated. Scalebars are 5 μm for full cells and and 1 μm for insets. A timelapse of reconstructed images for this cell are shown in Movie 4. **(B)** Immobile fraction of Ftractin-mEos3.2-GG evaluated over 30-second time intervals averaged across populations of cells that are unperturbed (N=15) or perturbed by pretreating with 1μM JSP (N=6) or expressing HLE (N=6). Values are normalized by the average immobile fraction over time intervals before clustering individual cells. Errors are shown as shaded regions and represent the standard error of the mean. **(C)** ⍵, ξ_0_ and v_exp_ determined from the autocorrelation of Ftractin-mEos3.2-GG evaluated over 2 or 1-min time intervals. Error represents the standard error of the mean for values determined from populations of cells that are untreated (N=12), perturbed with 1μM JSP (N=6), or expressing HLE (N=4). Significance testing is determined using a 1-tailed t-test. **(D)** Integrated intensity from the autocorrelation of BCR evaluated over 20-second time intervals and fit to *A*(1 − *exp*(− *t*/*t* )) + *C* for t>0 as described in Methods. **(E)** ⍵, ξ_0_ and v_exp_ determined from the autocorrelation of mEos3.2-Lifeact evaluated as described in C and averaged across a population of cells (N=6). Plots showing data points for individual cells for panels C and E appear in Figure S5. Symbols for p-values from significance testing represent *p≤0.05, **p≤0.01, ***p≤0.001, ****p≤0.0001, or ns for p>0.05

Focusing first on MP actin alone, we find that the immobile fraction of Ftractin-mEos3.2-GG rapidly decreases upon streptavidin addition **(Figure 5B)**, consistent with a reduction in MP actin density coincident with initial receptor clustering. After 1 min, MP actin density is replenished, recovering to pre-stimulation levels after roughly 4 mins. This transient reduction in MP actin is in good agreement with previous reports that have attributed actin disassembly to dephosphorylation of ezrin and activation of the actin-severing protein cofilin (37, 53–55). No transient reduction in the immobile fraction is detected in cells pretreated with JSP, which inhibits actin remodeling, or in cells expressing HLE, which has been previously indicated to inhibit signaling-dependent actin decoupling due to ezrin dephosphorylation (37).

Changes in average MP actin structure upon BCR clustering were also interrogated through the autocorrelation, g(r, *τ*), to extract the average steady state corral size (ξ_0_), the expansion speed (v_exp_), and remodeling rate (⍵) using the strategies described in Figure 3 but modified to be appropriate for single-color measurements (see methods) (**Figure 5C, Figure S5)**. Using this approach, we detect a transient increase in ⍵ over the first 2 minutes of BCR activation, consistent with the transient removal of MP actin from the membrane surface. A transient increase in ξ_0_ is also observed during the first minute after streptavidin addition, followed by a return to baseline values. We note that the values of ξ_0_ reported here are smaller than those reported in Figure 3, possibly because autocorrelation retains some correlations originating from single-molecule motions. This timing coincides with the loss of MP actin density as identified through the immobile fraction. In contrast, significant changes are not detected in v_exp_, indicating that lateral movements of the MP actin mesh are not meaningfully altered over the stimulation time-course. Analogous transient changes for ⍵ and ξ_0_ are not detected for any measured parameters in control cells pretreated with JSP or expressing HLE. Overall, these results indicate that streptavidin addition triggers a transient loss of MP actin and the opening of the meshwork, resulting in larger MP actin corals.

The actin remodeling described above occurs while BCR is assembling into clusters, so we next explored the dynamics of BCR cluster assembly in the same cells by tabulating autocorrelations of the BCR signal. **Figure 5D** shows how the integrated intensity of g(r, *τ*→0) (see methods), which is proportional to the average number of BCRs associated with clusters over time. This approach captures the rapid assembly of receptors into clusters upon the addition of streptavidin. The rise in the correlation intensity is fit to the exponential form *I*(*t* > 0*s*) = *A*(1 − *exp*(− *t*/*t*_0_)) , with an offset determined from the unstimulated values *I*(*t* > 0*s*), to extract the characteristic clustering time t_0_. Through this analysis, t_0_ is faster than the time resolution of the measurement (t_0_<20s) for untreated cells, and slows in cells pretreated with JSP or expressing HLE, with characteristic clustering times of t_0_ = 85 ± 21 s and 22 ± 7 s, respectively. These observations are consistent with previous studies exploring connections between actin structure in the assembly and trafficking of immunoreceptor clusters (49, 54–67). More specifically, our analysis suggests that MP actin remodels upon strepaviden addition to transiently increase the area of actin corrals, facilitating the rapid assembly of BCR into clusters.

Lastly, we compare the transient remodeling of MP actin observed by SM-MPAct with mEos3.2-Lifeact, a probe for global f-actin. In contrast to MP actin, the autocorrelation analysis detects only minor variations in the ⍵, ξ_0_, and v_exp_, parameters (**Figure 5E, Figure S5)**. This indicates that, on average, the larger f-actin network is not changing during early receptor clustering and activation, within the sensitivity and time resolution of this analysis. That signals are more robust when viewing SM-MPAct than lifeact indicates that the majority of changes are limited to f-actin in close proximity to the plasma membrane.

### Membrane-proximal actin colocalizes with B-cell receptors

Simultaneous detection of BCR and SM-MPAct localizations enables an analysis of MP actin in the vicinity of BCR during BCR clustering and activation. There are numerous examples where SM-MPAct trajectories stall in the vicinity of BCR clusters **(Figure 6A)**, suggesting that these regions are rich in MP actin. This impression is quantified by tabulating the crosscorrelation between BCR and immobile SM-MPAct for simultaneous observations, c(r,*τ*→0), as described in methods **(Figure 6B)**. Comparing c(r,*τ*→0) before and after clustering reveals that BCR weakly colocalizes with MP actin and that clustering BCR increases colocalization. This measurement is time-resolved by calculating c(r,*τ*→0) over smaller steady-state time windows, then summarized by reporting the amplitude of correlations (c(r<50nm,*τ*→0)), which represents the simultaneous enrichment of MP actin within BCR clusters. This time-course indicates that MP actin is recruited to BCR clusters over time, consistent with the expected recruitment and activation of the Arp2/3 complex at BCR signalosomes (51, 68, 69). MP actin enrichment at BCR clusters is not detected in cells pretreated with JSP or in cells expressing HLE **(Figure 6B)**, two conditions that also inhibited the transient drop in MP actin density upon streptavidin addition. These measurements suggest that this initial MP actin remodeling enables the reassembly of f-actin at BCR clusters downstream of receptor activation.

**Figure 6.**
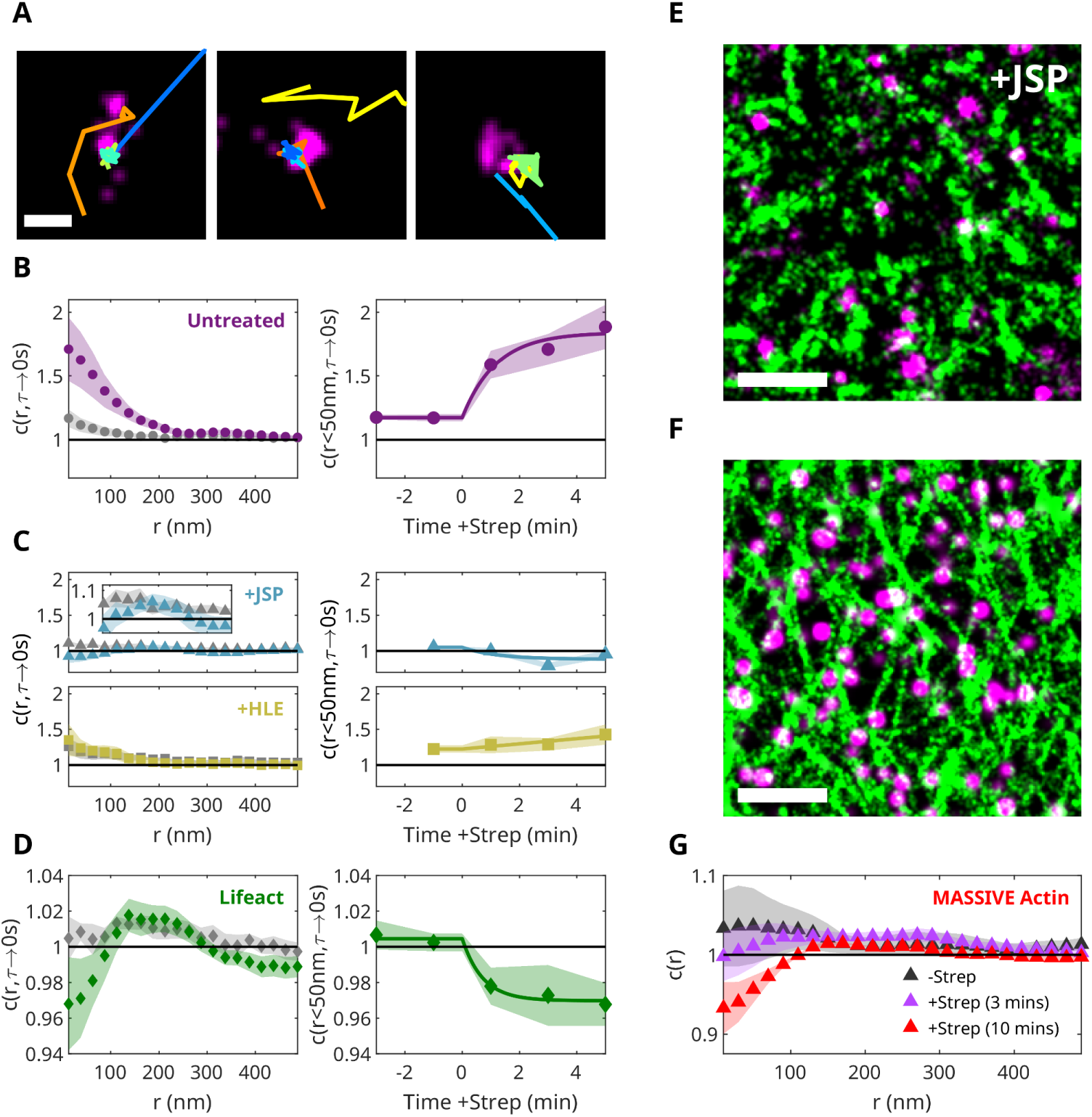
Membrane-proximal actin colocalizes with B-cell receptors. **(A)** Trajectories of Ftractin-mEos3.2-GG superimposed on reconstructions of BCR localizations acquired over 10s in cells treated with strepaviden. **(B)** *Left* - Simultaneous crosscorrelations (c(r,*τ*→0s)) between BCR and immobile Ftractin-mEos3.2-GG for localizations observed 1-minute before (-Strep) and 1-6 minutes after (+Strep) BCR clustering. *Right* - The amplitude of the simultaneous crosscorrelation (c(r<50nm, *τ*→0s)) between BCR and immobile Ftractin-mEos3.2-GG evaluated over 2-minute time windows, before and after BCR clustering. Plots depict the average correlation and standard error of the mean evaluated over a population of cells (N=15). **(C)** c(r,*τ*→0s) (left) and c(r<50nm,*τ*→0s) (right) between BCR and immobile Ftractin-mEos3.2-GG for cells pretreated with 1 μM JSP (N=6) or expressing HLE (N=6). The inset shows c(r,*τ*→0s) from r = 85 to 350 nm for the JSP-treated population with an expanded y axis to emphasize intensity at intermediate separation distances. **(D)** Super-resolution reconstruction of immobile Ftractin-mEos3.2-GG (green) and BCR (magenta) localizations accumulated 1-5 minutes after clustering for a representative cell pretreated with 1 μM JSP. **(E)** c(r,*τ*→0s) (left) and c(r<50nm,*τ*→0s) (right) between BCR and mEos3.2-Lifeact averaged across a population of cells (N=7). **(F)** Super-resolution reconstruction of MASSIVE actin (green) and BCR (magenta) for a representitive cell chemically fixed 10 minutes after BCR clustering. **(G)** c(r) between BCR and massive actin evaluated and averaged for cells fixed before (N=2), 3 minutes after (N=4), and 10 minutes after (N=2) BCR clustering. Scale bars in A, D, and F are 200 nm, 1 μm, and 1 μm, respectively.

Conducting the same analysis using localizations of BCR and soluble mEos3.2-lifeact reveals that BCR again tends to localize with total f-actin, but that this colocalization diminishes upon streptavidin addition (**Figure 6D**). This indicates that the total f-actin density is lower at the center of BCR clusters than the average density over the whole cell surface. Interestingly, elevated f-actin density is detected at separation distances of 100-200 nm, suggesting that BCR clusters reside near but not on f-actin filaments. This interpretation is supported by reconstructed images of BCR clusters and MP actin in cells pretreated with JSP, which inhibits actin assembly at BCR clusters **(Figure 6E)**, and by total f-actin in untreated cells imaged after chemical fixation (**Figure 6F**). Under both conditions, BCR clusters frequently reside beside regions of high actin density, and crosscorrelations detect elevated densities at separation distances of 100-200nm (**Figure 6C, G)**. The qualitative differences between observations with SM-MPAct and lifeact indicate that f-actin remodeling after strepaviden addition primarily occurs within the MP actin pool, which is a small fraction of the total f-actin detected under TIR. This MP actin pool includes both f-actin that contributes to the structural actin network and the f-actin specifically recruited to BCR signalosomes.

Our overall conclusions are summarized in the sketches of **Figure 7**. Initially, BCR is largely uniformly distributed in the plasma membrane. Immediately after streptavidin addition, MP actin rapidly remodels from the cell surface, effectively increasing the cortical meshwork size and enabling f-actin reassembly at BCR clusters. We speculate that opening the MP actin mesh enables more robust, efficient assembly of BCR clusters, as cluster assembly is slowed when opening is inhibited. The initial MP actin structure is restored within several minutes, and BCR clusters reside adjacent to cortical f-actin, suggesting that the structural actin plays a role in localizing BCR clusters. Our quantitative measurements provide evidence for a slow-moving structural actin network capable of regulating the organization of membrane components (43, 70–72) while also supporting the presence of more dynamic actin filaments that assemble at the plasma membrane in response to signaling processes and represent a minority of the total cortical f-actin (22, 23, 73, 74). This interpretation is in good agreement with past work showing that f-actin plays dual roles: restricting the mobility of BCR (54, 75) and regulating the motion and functions of BCR clusters downstream of BCR activation (56, 58).

**Figure 7.**
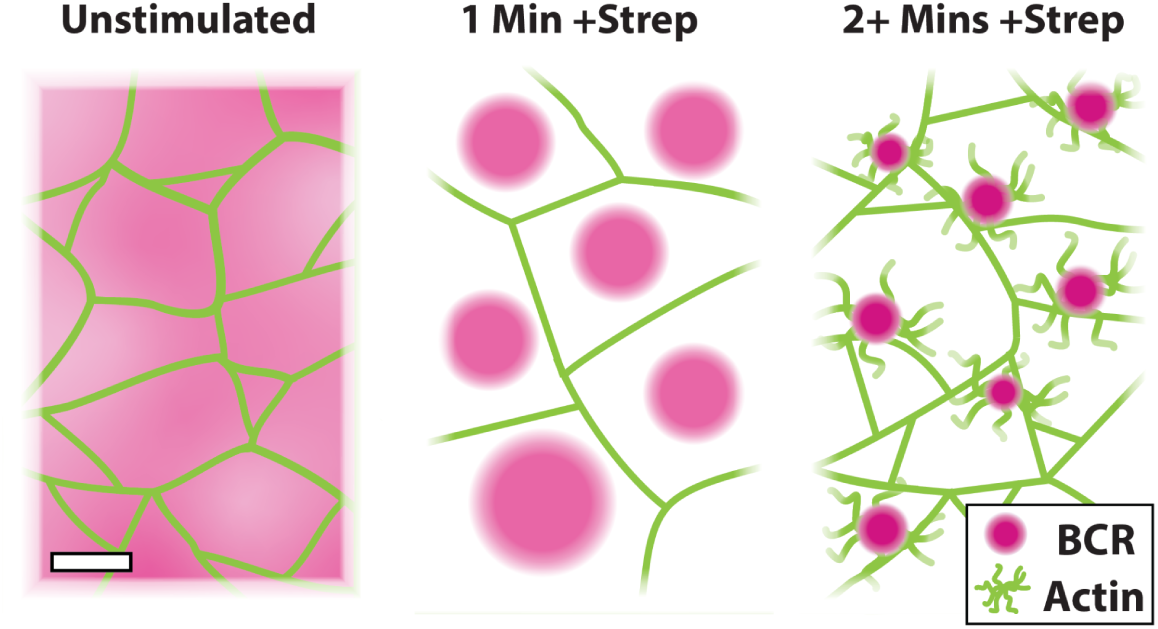
A model for MP actin remodeling during early BCR activation. Actin remodeling during BCR activation enables rapid clustering of BCR and assembly of MP actin at BCR cluster sites. Left - Time-averaged distribution of BCR (magenta) on a membrane compartmentalized by an MP actin meshwork (green) in the unstimulated state. The scale bar symbolizes 200 nm. Center - Assembly of BCR clusters and simultaneous remodeling of MP actin during the first minute after BCR cross-linking, leading to larger actin corals. Right - MP actin enrichment at BCR cluster sites and compartmentalization of BCR clusters by the recovered cortical actin mesh.

## CONCLUSIONS

SM-MPAct is a single-molecule probe embedded in the plasma membrane that becomes transiently immobilized through interactions with MP actin. The positions of these binding events can be identified by monitoring probe mobility, and the fraction of bound probes at any given time reflects the average density of MP actin across the cell surface. While spatial sampling is too slow to visualize dynamic MP actin structures directly in unperturbed cells, this approach can resolve changes in the average MP actin density on the order of seconds in cells treated with cytoskeleton-altering drugs or undergoing signaling responses.

Through transient binding, SM-MPAct localizations sample the underlying MP-actin structure in both space and time. The motions of SM-MPAct probes not bound to actin are largely uncorrelated with each other, due to the relative homogeneity of the cell surface. In contrast, probes bound to f-actin remain correlated, even over long times, because they repeatedly sample the same spatially heterogeneous MP actin structures. Recognizing this, we developed an approach using correlation functions to measure average properties of MP actin in live cells, including the size of the MP actin mesh, its expansion speed, and its turnover rate. We find that the meshwork size determined in live cells agrees well with fixed-cell measurements, that dynamic properties agree well with literature values obtained using other experimental approaches, and that chemical and physical perturbations of actin and actin-binding proteins produce the expected changes in MP actin structure and dynamics (38, 42, 43, 46). Importantly, these same methods can be applied to cells undergoing signaling responses, reporting on the changes in actin structure and dynamics that contribute to cell signaling processes.

Overall, SM-MPAct is sensitive to the pool of f-actin that directly interacts with plasma membrane components, and the analytical tools we have developed have the sensitivity and temporal resolution to detect changes in MP actin structure and dynamics relevant to signaling processes. While we have focused primarily on MP actin in B cells and its changes during BCR signaling, we anticipate that the experimental and analytical tools developed could be extended to other cell systems and processes in which actin structure and dynamics are known to play important roles, such as endocytosis, cell migration, and synapse formation (1).

## AUTHOR CONTRIBUTIONS

S.L.V. devised the central questions of the study. A.D. and S.L.V. developed the experimental methods. A.D. carried out all of the experiments. A.D. and S.L.V. analyzed the experimental data. A.D. and S.L.V. wrote the manuscipt.

## ACKNOWLEDGEMENTS

We thank Jennifer Flanagan-Natoli for performing preliminary measurements, Andrea Stoddard for assistance with generating SM-MPAct constructs, and Thomas Shaw, Sarah Shelby, Natalie Rogers, and Yousef Bagheri for helpful conversations. Research was supported by the NIH (R35GM152150 to SLV).

## MATERIALS AND METHODS

### Constructs

DNA sequences for SM-MPAct constructs were based on previously described membrane probes (40) modified by attaching the Lifeact or Ftractin actin-binding domains to the N or C-terminus, respectively. Ftractin-mEos3.2-GG, Ftractin-mEos3.2-FP, and Ftractin-mEos3.2-FPP were constructed by adding the sequence of Ftractin and its linking amino acids (20) to the N-terminus of the mEos3.2-GG, mEos3.2-FP, and mEos3.2-FPP probes (40). Similarly, the PM-mEos3.2-Lifeact and M-mEos3.2-Lifeact were constructed by adding the sequence of Lifeact and its linking amino acids (20) to the C-terminus of PM-mEos3.2 and M-mEos3.2 (40). PM-Halo-Lifeact was constructed by adding a Halo tag and a short chain of linking amino acids in place of mEos3.2 in the PM-mEos3.2-Lifeact probe. The FtractinΔ1-6-mEos3.2-GG mutant was modified from the Ftractin-mEos3.2-GG by removing the 6 N-terminal amino acids (20). Lifeact-mEos3.2 was obtained from Barbara Baird at Cornell University (76). Amino acid sequences for SM-MPAct probes are provided in **Table S1**. The membrane-bound constitutively active ezrin mutant (HLE) was obtained from Neetu Gupta at the Cleveland Clinic (37).

### Cell Culture and Sample Preparation

CH27 cells were cultured as described previously (39), and transfected approximately 24 hours prior to imaging using the Lonza electroporation system with the SF nucleofactor kit and program CA-137. Cells were plated on MatTek dishes coated with VCAM-1, prepared as described previously (40), for 15-20 minutes before thoroughly washing. U2OS cells were cultured as described previously (77) and transfected 24 hours before imaging with the Lonza SE nufleofactor kit and program CM-104. Cells transfected with Halo-tag constructs were incubated with 1 nM of the JF-635b (27) halo-ligand, acquired from Luke Lavis (Janelia Research Campus), for one hour in cell media before plating. For BCR-clustering experiments, CH27 cells were labeled for 10 min after plating with 5 μg mL^−1^ Goat ɑMouse IgM Fab (μ-chain specific) conjugated with biotin and SiR (Fab-biotin-SiR), prepared as described previously (39, 40).

For fixed cell experiments, CH27 cells were plated on VCAM-1-coated dishes, prepared using a Rabbit ɑHuman IgG antibody in place of the Goat ɑHuman IgG, and labeled with a commercial biotin-conjugated Goat ɑMouse IgM Fab (μ-chain specific) (Jackson ImmunoResearch 115-067-020). BCR was cross-linked with 1 μg mL^−1^ AF488-labeled streptavidin for varying amounts of time before fixation with 4% PFA in 1X PBS. After 10 minutes of fixation, samples were blocked and permeabilized with blocking solution (2% w\v BSA, 0.01% Triton X-100 in 1X PBS). BCR clustering was confirmed visually through excitation at 488nm. Fixed samples were then further labeled with 25 μg mL^−1^ ɑGoat IgG conjugated with a DNA docking strand (Purchased from Massive Photonics) and imaged in Massive Photonics imaging buffer with the complementary DNA imaging strand conjugated to Atto655, as well as the MASSIVE photonics f-actin PAINT reagent conjugated to Cy3b. Concentrations of PAINT imaging reagents were adjusted for the desired density of localizations.

### Single-Molecule Imaging

For most live cell experiments, cells were imaged in a live cell-compatible buffer (20 mM hepes, 5.6 mM glucose, 135 mM NaCl, 1.8 mM CaCl_2_, 5 mM KCl, 1 mM MgCl_2_, 0.2% w\v BSA, 40 μg mL^−1^ catalase, pH 7.4). Measurements involving BCR localization were conducted in a reducing environment to enable photoswitching of SiR (30 mM Tris, 100 mM NaCl, 5 mM KCl, 1 mM MgCl_2_, 55.6 mM Glucose, 1.8 mM CaCl_2_, 9.8 mM glutathione, 200 μg mL^−1^ catalase, 100 μg mL^−1^ glucose oxidase, pH 8.0).

Live samples were imaged using an Olympus IX81 inverted microscope with a 100X objective lens (NA 1.49) and ZDC autofocus. Samples expressing mEos3.2 constructs were excited with a Coherent LS 561 nm 120 mW fiber pigtail laser at 10 mW and a Coherent CUBE 405 nm 50 mW fiber pigtail laser at 0.13-0.39 mW, varied to obtain an optimal density of single molecule localizations. Samples that were labeled with JF635b halo-ligand or Fab-biotin-SiR were excited with a Coherent OBIS LX 647 nm 100mW fiber pigtail laser at 100 mW. TIRF microscopy was enabled with a cellTIRF module. For 2-color experiments, fluorescence emission was filtered using a Photometrics DV2 channel splitter equipped with a T640lpxr dichroic mirror as well as a 585/40 nm band pass filter to observe near-red emission and a 700/75 nm band pass filter or a 685 nm long pass filter to observe far-red emission for imaging JF635b or SiR, respectively. An Andor iXon Ultra 897 EMCCD camera was used to record the fluorescence emission of samples with a constant acquisition time of 20 ms per frame for 1-color experiments or 30 ms per frame for 2-color experiments. The actual framerate includes the time delay that occurs from the reading of the EMCCD camera. Cells were imaged for 10,000-15,000 frames. For temperature modulations above or below room temperature (23 °C), cells were imaged in a KRi sample temperature feedback cooling & heating stage top Incubator system equilibrated to the indicated temperature before imaging and maintained within ± 2 °C of the indicated temperature throughout imaging. The temperature of samples was detected with a Tokai-Hit bendable wire thermal probe placed directly into sample dishes.

Treatment with pharmacological inhibitors included 0.5 μM latrunculin A (Cayman 10010630), 1 μM jasplakinolide (Cayman 11705), 100 μM CK666 (Sigma SML0006), 40 μM SMIFH2 (Sigma 344092), 25 μM para-amino-blebbistatin (Cayman 22699), or 5 μM pentachloropseudilin (SML3996). All inhibitors were diluted from DMSO stock solutions, except for LatA, which was stored in an ethanol stock solution. Inhibitor (or 2% (v/v) DMSO vehicle solution) was added directly to dishes, either before imaging or throughout imaging, where indicated. BCR labeled with Fab-biotin-SiR was cross-linked with 1 μg mL^−1^ streptavidin. For experiments involving observation of the same cell before and after cross-linking or treatment inhibitors, cells were imaged for 10,000-15,000 frames, both before and after treatment.

For fixed cell imaging, samples were imaged using an Olympus IX83-XDC inverted microscope with a 100X UAPO TIRF objective lens (NA 1.49) and ZDC autofocus. Samples were excited with a Coherent Sapphire 561 nm LP laser and an OBIS LX 647 nm 120 mW laser. Fluorescence emission was filtered through a Hamamatsu w-view gemini splitter equipped with 575/40 nm band pass and 685 nm long pass filters, to observe near-red and far-red emission, respectively. An Andor iXon 897 EMCCD camera was used to record the fluorescence emission of samples with a constant acquisition time of 100 ms per frame. Cells were imaged for 10,000-15,000 frames.

### Image Processing and Single Particle Tracking

Single molecules were localized using custom MATLAB software that implements maximum-likelihood fitting for single emitters (78), or the Thunderstorm Fiji plugin with multi-emitter fitting for experiments with SiR or PAINT probes (79). Localization errors are returned directly from the fit as the Cramér-Rao lower bound for the maximum-likelihood estimate. Transforms for 2-color data and drift correction were applied to localized positions, as described previously (41, 80). When appropriate, time intervals (τ) were corrected to account for the finite integration window according to τ_*c*_ = *frame time* * (1 − (1/3)(*integration time*/τ)) (81).

Single molecules were tracked with custom MATLAB software that links localizations across sequential frames within 500 nm trajectory steps, terminating trajectories at points with more than one localization pair. Tracking data is further analyzed with SASPT (26), which initially splits trajectories into segments with at most 10 steps (11 frames) per segment. The SASPT code produces a 2-dimensional (2D) probability distribution describing localization error (σ) and diffusivity (D). This 2D distribution is collapsed onto one dimension by summing over σ to produce the distributions shown, resulting in diffusion coefficients that can be smaller than those expected due to localization error alone. The posterior distribution was integrated below D ≤ 0.01 μm² s^−1^ to get the fraction of immobile probes. The mobility of individual track segments were assigned by selecting the peak of the 1D posterior distribution for each segment. For time-resolved measurements, the SASPT posterior distribution was calculated over 100 frame intervals (2-3 s).

### Correlation Function Analysis and Fitting

Spatiotemporal correlation functions were calculated from masked localizations with a gradient correction, as described previously (39, 41). Briefly, the distances between pairs of localizations were assembled into histograms with specified spatial and temporal bins, and then normalized to account for the finite spatial and temporal extent of the dataset. These functions were additionally normalized to eliminate long-range spatial and temporal gradients inherent to the dataset (e.g. illumination intensity gradients in space and photobleaching in time). While spatial bins were always uniformly spaced, temporal bins were sometimes constructed with different widths to improve statistics at larger *τ*, where the finite extent of the temporal window results in fewer probe pairs per frame. Errors in values for individual cells are estimated from the number of pairs that contribute to the measurement, normalized by the same factors used to normalize the cross-correlation. Typically, reported errors represent the standard error of the (unweighted) mean across multiple cells.

When noted, we tabulate the integrated intensity of either auto- or cross-correlations as *I*(*τ*) = ∑ 2π*r*(*g*(*r*, τ) − 1)Δ*r*, for r < 600 nm, where *r* is the spatial bin center and Δ*r* is the spatial bin width. Where noted, we estimate simultaneous crosscorrelations, *c*(*r*, *τ* → 0), by averaging *c*(*r*, *τ*) between 0. 2*s* < *τ* < 2*s* to improve statistics and avoid contributions from bleed-through between imaging channels. Also where noted, we estimate the simultaneous autocorrelation (*g*(*r*, *τ* → 0)) by averaging *g*(*r*, *τ*) between 0. 1*s* < *τ* < 0. 2*s* to reduce the impact of overcounting of SiR fluorophores for *τ* < 0. 1*s*.

Both autocorrelations, *g*(*r*, τ), and crosscorrelations, *c*(*r*, τ), were parameterized by fitting to functional forms as described throughout the manuscript. Fitting was accomplished using the fit() function in Matlab, and was typically weighted using the inverse variance of fitted datapoints. Errors in fit parameters are 68% confidence intervals determined using the confint() function in Matlab.

The mobility of SM-MPAct probes were determined from fitting the *g*(*r*, τ) at constant τ to a sum of two Gaussians, representing slow and fast mobility states (39–41): 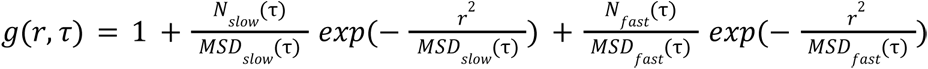 where *N*_*slow*_ and *N*_*fast*_ are proportional to the number of slow and fast correlated molecules, and *MSD*_*slow*_ and *MSD*_*fast*_ is the mean squared-displacement of the slow and fast populations evaluated at every time-interval. The actin-binding time, *τ*_off_, for SM-MPAct is determined by fitting the fraction of probes in the slow fraction *N*_*slow*_ (τ)/(*N*_*slow*_ (τ) + *N*_*fast*_ (τ)) to *A exp*(− *τ*/*τ*_*off*_) + *C*.

The turnover rate, ω, was determined by fitting *c*(*r*, τ) or *g*(*r*, τ) over at least 14 *τ* -intervals spanning τ = 1-70s for *c*(*r*, τ) or τ = 5-60s for *g*(*r*, τ). When the measurement was time-resolved, correlation functions were tabulated using 120s of acquired localizations in each time-window, otherwise the whole dataset was used. The correlation intensity was then fit to *I*(*τ*) = *A exp*(− ω*τ*) + *C*. When appropriate, bounds were included on some fit parameters to reduce poor fits due to noise, especially when the use of inhibitors led to slowly decaying curves. The characteristic meshsize, ξ_0_, and the expansion speed, *v*_*exp*_, were determined by evaluating *c*(*r*, τ) or *g*(*r*, τ) for at least 11 logarithmically spaced *τ*-intervals between τ = 1-30s for *c*(*r*, τ) or τ = 5-30s for *g*(*r*, τ). When the measurement was time-resolved, correlation functions were tabulated using 60s of acquired localizations in each time-window, otherwise the whole dataset was used. Raw correlation functions were then fit to *c*(*r*, *τ*) = 1 + *A*/(2πξ(*τ*) ) *exp*(− *r*/ξ(*τ*)) + *C*, where ξ(*τ*) represents the decay length of the correlation. ξ(τ) was then further parameterized by fitting to ξ(*τ*) = ξ_0_ + *v*_*exp*_ *τ*. Cells with a small number of pairs due to low expression were removed from the dataset used for time-resolved measurements.

The characteristic time for BCR clustering, *t*_0_, was determined by tabulating the *g*(*r*, τ) over 20s time-windows both before and after BCR crosslinking with streptavidin. The time-resolved intensity of BCR autocorrelations was then fit to *I*(*t* ≥ 0*s*) = < *I*(*t* < 0*s*) >+ *A*(1 − *exp*(− *t*/*t*_0_)).

## MOVIE CAPTIONS

**Movie 1: Single-molecule microscopy of SM-MPAct.** Raw fluorescence images of Ftractin-mEos3.2-GG expressed in a CH27 cell and imaged under TIR excitation. Images were acquired at approximately 20 frames per second. Scale bar is 5 μm.

**Movie 2: Single-molecule microscopy of SM-MPAct and non-actin binding membrane probes.** Raw Fluorescence images of Ftractin-mEos3.2-GG (left), mEos3.2-GG (middle), and FtractinΔ1-6-mEos3.2-GG (right) expressed in different CH27 cells imaged under TIR excitation. Images were acquired over varying acquisition times, ranging from 20 to 30 frames per second, as displayed. Scale bar is 5 μm.

**Movie 3: Time-lapse of immobile SM-MPAct in a CH27 cell.** Movie composed of reconstructed images from immoble Ftractin-mEos3.2-GG localizations from the representative cell in Fig. 4C. Each image is reconstructed from localizations acquired over 1000 frames (approximately 23s), which is near the remodeling rate (⍵) for this cell, incremented by 150 frames (approximately 3.3s). The movie is rendered over the entire experimental time, with the elapsed time shown in the top right corner. Scale bar is 5 μm.

**Movie 4:Time-lapse of BCR and MP actin before and after BCR clustering.** Movie composed of reconstructed images from immobile Ftractin-mEos3.2-GG (green) and BCR (magenta) localizations from the representative cell in Fig. 5A. Each image is reconstructed from localizations observed over 1000 frames (approximately 30s), incremented by 150 frames (approximately 4.8s). The movie depicts the entire experimental time before and after clustering, with the real elapsed time shown in the top right corner. The time point “0:00” represents the first time point immediately after the addition of streptavidin. Scale bar is 5 μm.

