## Supplemental for "A single-molecule reporter of membrane-proximal actin detects rapid remodeling upon B cell receptor clustering"

| Plasmid Name (N to C) and Description | Amino Acid Sequence (N to C) |
|---|---|
| Ftractin-mEos3.2-GG:<br>Geranylgeranylated sequence from KRas GTPase linked to mEos3.2 and Ftractin with a 7 AA linker. | Ftractin-SDPPVAT-mEos3.2-FRSDGKKKKKKSKTKCQLL |
| Ftractin-mEos3.2-FPP:<br>Peptide sequence from HRas containing 1 farnesylation and 2 palmitoylations linked to mEos3.2 linked to Ftractin with a 7 AA linker. | Ftractin-SDPPVAT-mEos3.2-FSSSSLNSGCMSCKCVLS |
| Ftractin-mEos3.2-FP:<br>Peptide sequence from NRas containing 1 farnesylation and 1 palmitoylation linked to mEos3.2 linked to Ftractin with a 7 AA linker. | Ftractin-SDPPVAT-mEos3.2-FSSSSLNSAVDGCMGLPCVVM |
| M-mEos3.2-Lifeact:<br>Myristoylated sequence from Src15 linked to mEos3.2 linked to Lifeact with a 5 AA linker. | MGSSKSKPKDPSQRRNNNNGPVAT-mEos3.2-SAGGT-Lifeact |
| PM-mEos3.2-Lifeact:<br>Palmitoylated and miristoylated sequence from Lyn linked to mEos3.2 and Lifeact with a 5 AA linker. | MGCIKSKRKDKDLELKLRILQSTVPRARDPPVAT-mEos3.2-SAGGT-Lifeact |
| PM-Halo-Lifeact<br>PM-mEos3.2 (1) sequence modified with a halo tag and linked to Lifeact with a 5 AA linker. | MGCIKSKRKDKDLEPTTDNDGS-Halotag-SAGGT-Lifeact |
| FtractinΔ1-6-mEos3.2-GG:<br>Truncated non-actin binding mutant of GG-mEos-Ftractin with a 7 AA linker. | FtractinΔ1-6-SDPPVAT-mEos3.2-FRSDGKKKKKKSKTKCQLL |

**Supplemental Table 1: Sequence and design of SM-MPAct probes.** Descriptions and sequences are provided for the SM-MPAct probes used in this study. Sequences for GG, FPP, FP, M, and PM are described previously (40). Underlined portions represent the sequences taken from native proteins. Sequences for Ftractin (full-length and truncated mutant), Lifeact, mEos3.2, and the Halo tag are omitted from the table but listed below.

mEos3.2 (40):
MSAIKPDMKIKLRMEGNVNGHHFVIDGDGTGKPFEGKQSMDLEVKEGGPLPFAFDILTTAFHYGNRVFAKYPDNIQDYFKQSFPKGYSWE
RSLTFEDGGICNARNDITMEGDTFYNKVRFYGTNFPANGPVMQKKTLKWEPSTEKMYVRDGVLTGDIEMALLLEGNAHYRCDFRTTYKA
KEKGVKLPGAHFVDHCIEILSHDKDYNKVKLYEHAVAHSGLPDNARR

Halo Tag:
EIGTGFPFDPHYVEVLGERMHYVDVGPRDGTPVLFLHGNPTSSYVWRNIIPHVAPTHRCIAPDLIGMGKSDKPDLGYFFDDHVRFMDAFI
EALGLEEVVLVIHDWGSALGFHWAKRNPERVKGIAFMEFIRPIPTWDEWPEFARETFQAFRTTDVGRKLIIDQNVFIEGTLPMGVVRPLTEV
EMDHYREPFLNPVDREPLWRFPNELPIAGEPANIVALVEEYMDWLHQSPVPKLLFWGTPGVLIPPAEAARLAKSLPNCKAVDIGPGLNLLQ
EDNPDLIGSEIARWLSTLEISG

Ftractin (20): MGMARPRGAGPCSPGLERAPRRSVGELRLLFEARCAAVAAAAAAG

FtractinΔ1-6 (20): MGAGPCSPGLERAPRRSVGELRLLFEARCAAVAAAAAAG

Lifeact (20): MGVADLIKKFESISKEE

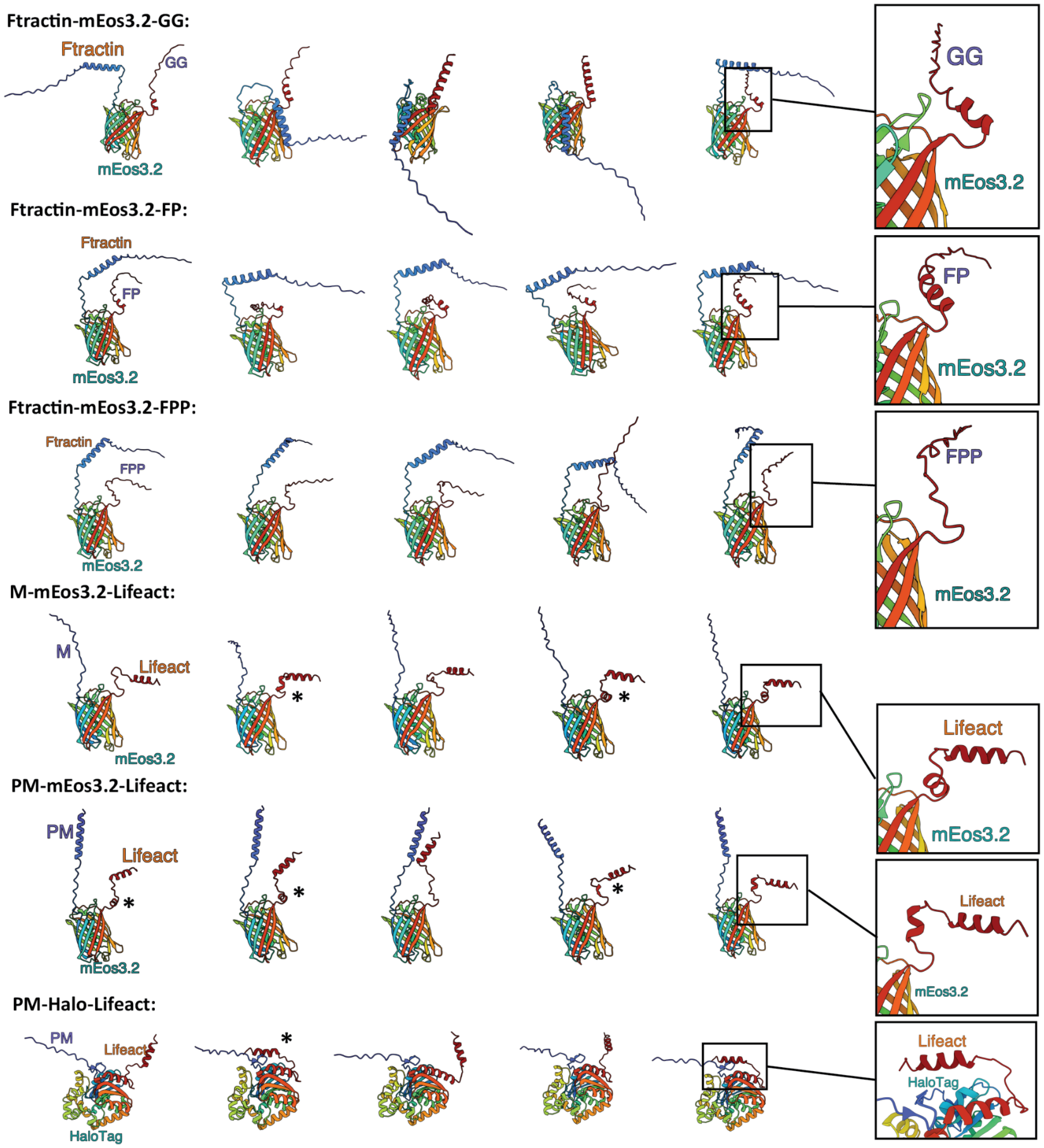


**Figure S1. Alpha fold predictions reveal differences in the secondary structure and conformations of SM-MPAct probes.** AlphaFold3 (28) structural predictions from five randomly generated seeds for the SM-MPAct probes used in this study. Insets highlight important structural features and conformations that differ across comparable probes. For example, Ftractin-mEos3.2-GG and Ftractin-mEos3.2-FP contain an alpha helix in the linking sequence between mEos3.2 and the membrane-binding motif that is unstructured in Ftractin-mEos3.2-FPP. Additionally, several predicted structures for M-mEos3.2-Lifeact (3/5) and PM-mEos3.2-Lifeact (4/5) contain an alpha helix in the linking sequence between mEos3.2 and Lifeact. It is possible that these differences in secondary structure could impact the ensemble of orientations available to the f-actin binding sites of these probes, giving rise to the differences detected in their immobile fractions shown in Figure 1F. More strikingly, several structures of PM-Halo-Lifeact (2/5) suggest the presence of specific interactions between lifeact f-actin binding domain and the halotag surface. This apparent competition for Lifeact binding may account for the greatly reduced immobile fraction observed for this probe as shown in Figure 1F. All renderings and zoom-ins were made using the interactive AlphaFold Server.

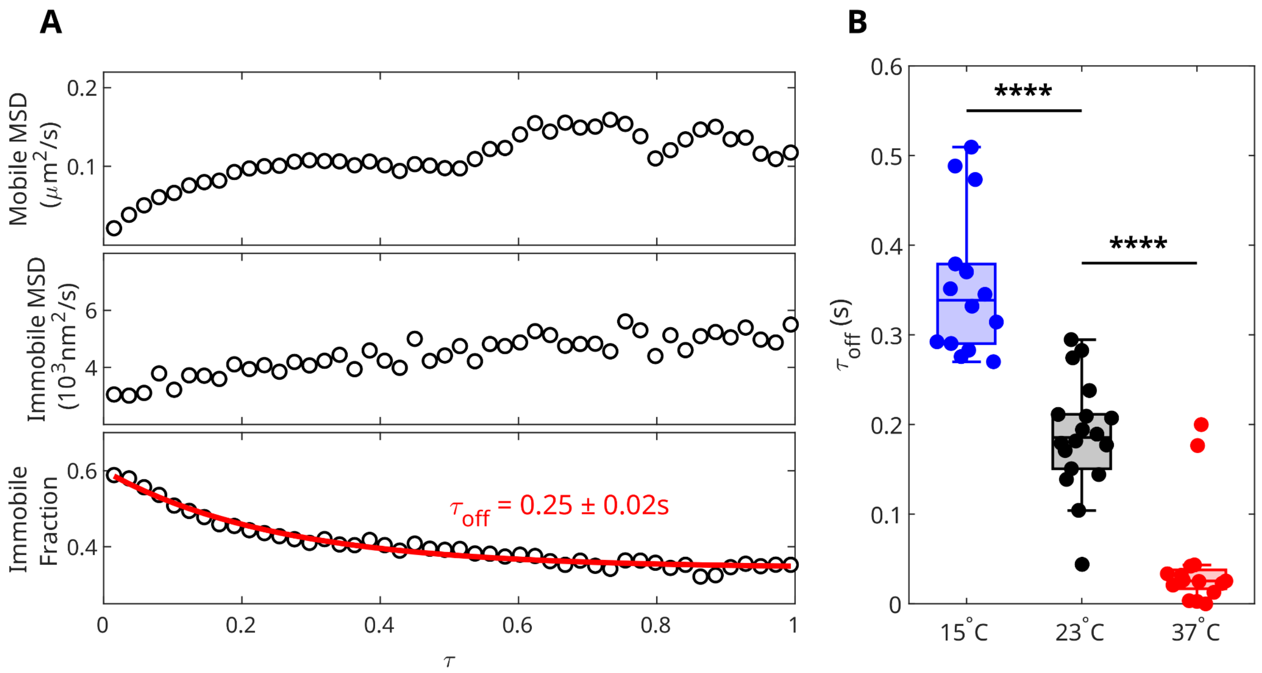


**Figure S2. Determination of the actin off-time for SM-MPAct. (A)** Mobile and immobile populations of Ftractin-mEos3.2-GG can be identified through fitting the autocorrelation at each $\tau$ to $g(r) = 1 + N_{immobile}/MSD_{immobile}\, exp(-r^2/MSD_{immobile}) + N_{mobile}/MSD_{mobile}\, exp(-r^2/MSD_{mobile})$. Here, fit parameters are shown for the representative cell observed at 23°C in Figure 3A. $N_{immoble}$ and $N_{mobile}$ are proportional to the number of immobile and mobile molecules, respectively. The immobile fraction is defined as $F_{immobile} = N_{immobile}/(N_{immobile} + N_{mobile})$. $F_{immobile}$ decays with time as probes unbind from f-actin and enter the mobile pool. This off-time, $\tau_{off}$, is evaluated by fitting $F_{immobile}$ to $A\, exp(-\tau/\tau_{off}) + C$. $\tau$ is corrected for the integration time as described in the methods. **(B)** $\tau_{off}$ for PM-mEos3.2-Lifeact observed across populations of cells imaged at 15°C (N=14), 23°C (N=18), and 37°C (N=16). Statistical significance is determined with a two-tailed t-test. Symbols for p-values from significance testing represent *p≤0.05, **p≤0.01, ***p≤0.001, ****p≤0.0001, or ns for p>0.05

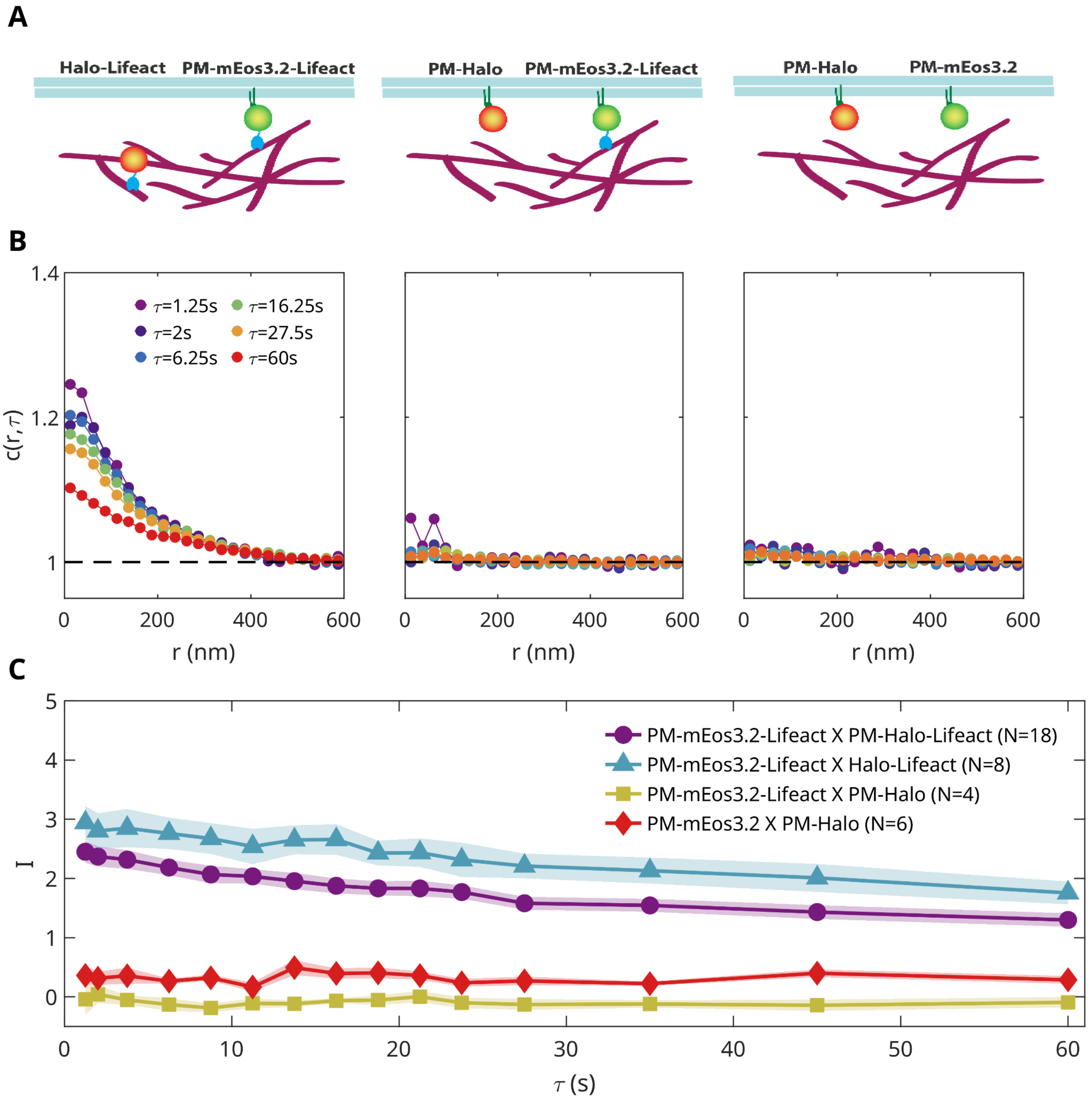


**Figure S3: Two-color long-$\tau$ cross-correlations of actin and membrane-binding probes. (A)** Cartoons depicting experiments for cross-correlation comparisons in panel B. **(B)** Example two-color cross correlations between PM-mEos3.2-Lifeact and Halo-Lifeact (left), PM-mEos3.2-Lifeact and PM-Halo (middle), or PM-mEos3.2 and PM-Halo (right), evaluated between $\tau$=0.1-70s. **(C)** Integrated intensity of all cross-correlation comparisons averaged across populations of cells. Error represents the standard error of the mean across cells.

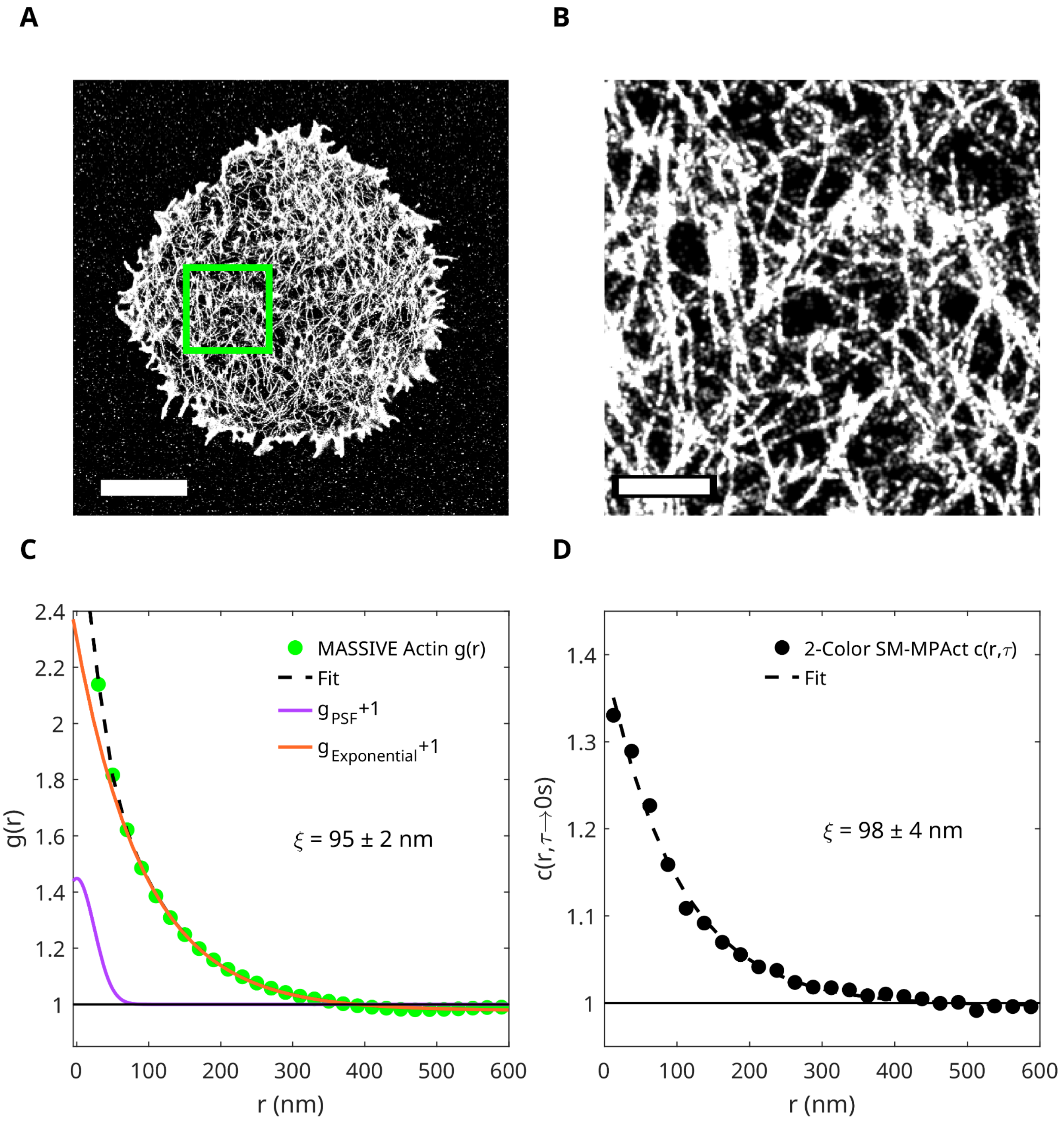


**Figure S4: Simultaneous SM-MPAct crosscorrelations from live cells resemble autocorrelations of total actin detected in chemically fixed cells. (A)** Reconstructed image of MASSIVE actin localizations acquired through imaging of a chemically fixed CH27 cell. This probe labels total f-actin. Scale bar is 5 µm. **(B)** Zoom-in of global actin reconstruction in panel A. Scale bar is 1 µm. **(C)** Autocorrelation, g(r), from localizations shown in panel A. g(r) is fit to $1 + A\,exp(-r/\xi) + Bexp(-r^2/(4\pi\sigma^2))$. The first (exponential) term captures the spatial decay of f-actin correlations, while the second (Gaussian) term accounts for over-counting of fluorophores. The exponential and Gaussian portions of the function are plotted together and separately. **(D)** Crosscorrelation between PM-mEos3.2-Lifeact and PM-Halo-Lifeact extrapolated to simultaneous observations ($c(r, \tau \rightarrow 0)$) as described in Methods for a representative cell. $c(r, \tau \rightarrow 0)$ is fit to $1 + Aexp(-r/\xi) + C$.

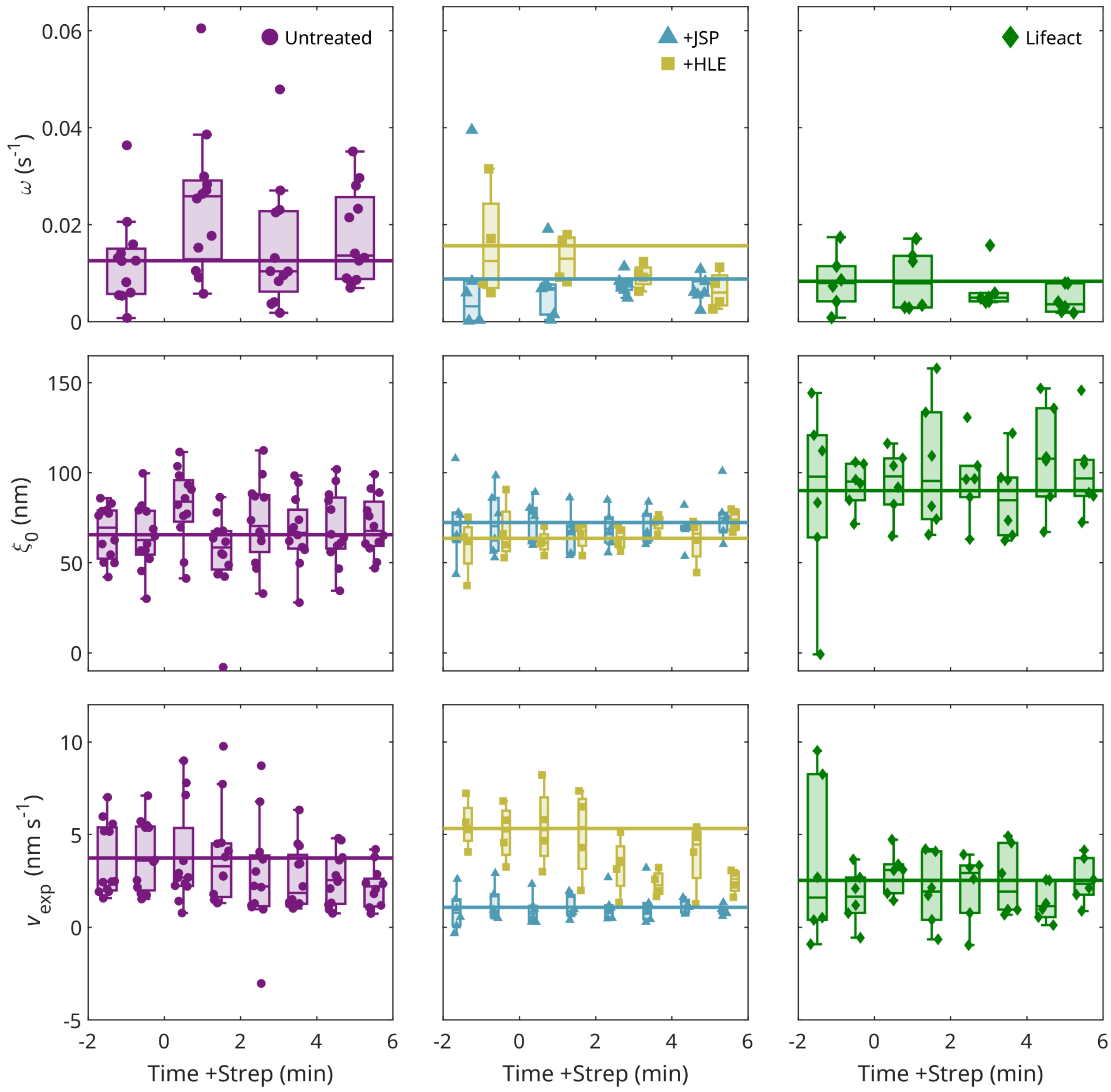


**Figure S5: Extended data of autocorrelation values quantified during BCR clustering.** $\omega$,$\xi_0$, and $v_{exp}$ values for individual cells for the time-resolved conditions shown in Figures 5C,E of the main text. For all measurements, N=12, N=6, N=4, and N=6 cells were evaluated for untreated, JSP-treated, HLE-expressing, and mEos3.2-Lifeact-expressing populations, respectively. Horizontal lines represent the average of the fit parameter for windows <0 min.
